# Cell-resolved transcriptional responses during heat-induced coral bleaching and recovery

**DOI:** 10.64898/2026.07.18.739195

**Authors:** Xavier Grau-Bové, Laia Montes-Espuña, Ewa Księżopolska, Sebastián R. Najle, Tali Mass, Shani Levy, Arnau Sebé-Pedrós

## Abstract

Coral reefs depend on an intracellular symbiosis between cnidarian hosts and photosynthetic dinoflagellates that is disrupted by ocean warming, causing coral bleaching. Yet how these responses are distributed across coral cell types during stress and recovery remains unclear. Here, we generate a temporal single-cell transcriptomic atlas of the reef-building coral Stylophora pistillata, profiling 20,000 cells during seven days at 32 °C and after 45 days of recovery. Heat stress reduced symbiont abundance and photosynthetic efficiency, followed by partial restoration. Early oxidative-stress, detoxification, and proteostasis programmes gave way to mitochondrial, cytoskeletal, and vesicular remodelling. These shared responses accompanied cell type-specific changes: epidermal cells activated tissue-renewal programmes, calicoblasts repressed calcification genes, and alga-hosting cells progressively lost transport, lipid-biosynthesis, and nutrient-exchange functions. Targeted profiling of sorted alga-positive cells revealed rare transcriptional states, including a population consistent with symbiont engulfment. Our results reveal how bleaching and recovery are coordinated across coral tissues and symbiotic cell states.

## Introduction

Coral reefs are among the most diverse ecosystems in the ocean, sustaining complex food webs and essential ecological functions, while providing habitat for roughly a quarter of all marine species^1–4^. Yet reefs are declining rapidly, with global living coral cover reduced by about 50% world - wide^2^. This loss is linked to ocean warming and to the increasing frequency, duration, and intensity of marine heat waves^5,6^.

A primary mechanism underlying reef degradation is heat-induced breakdown of coral algal symbiosis. Most reef-building corals host photosynthetic dinoflagellates of the Symbiodiniaceae family within specialised gastrodermal cells, where the host regulates symbiont abundance and physiology^1,7,8^. This intracellular partnership enables corals to thrive in oligotrophic waters: algal symbionts fix carbon and transfer a substantial fraction of this photosynthate to the host, receiving inorganic nutrients and a protected cellular niche in return^1,9^. Elevated temperatures can disrupt this relationship by impairing symbiont photophysiology and increasing oxidative stress, which initiates host cellular stress responses and can lead to symbiont loss, a process commonly termed as coral bleaching^10,11^. Bleaching rapidly reduces coral physiological performance, as sym-biont-derived photosynthates often supply over 90% of its energy requirements. In many species, the resulting deficit often results in coral mortality^12–14^.

Four global mass coral bleaching events of increasing extent and severity have been recognised. The latest, which began in 2023, exposed approximately 84% of the world’s reefs to bleach-ing-level heat stress^15–23^. In the summer of 2024, a severe marine heat wave triggered the first documented bleaching in the Gulf of Aqaba (Red Sea). Corals in this region tend to be thermally resilient, with common species able to tolerate 31 °C without severe bleaching^24–27^. Among them, *Stylophora pistillata*, stands out as one of the most abundant reef-building corals in the Gulf of Aqaba and a broadly distributed Indo-Pacific species. *S. pistillata* shows little to no bleaching between 31 °C to 32 °C but reaches a physiological tipping point beyond 33 °C, when bleaching, tissue loss and strong metabolic costs rapidly accrue^24,28–30^.

Bulk transcriptomic studies have identified conserved coral responses to heat stress, including the induction of chaperones, oxidative stress defences, shifts in apoptosis, immune signalling, and growth-related processes^31–34^. However, whole-organism measurements average signals over a mosaic of cell types, which may mask specificities in the timing and nature of the stress response expression programmes, vulnerable states, and recovery dynamics. As a result, it remains unclear which cell types initiate stress responses, how different these responses are between cells, how symbiosis-related expression programmes change during heat exposure, and to what extent distinct cell populations return to baseline after recovery. Single-cell transcriptomics can resolve these processes at a cellular resolution, and has already proved effective to investigate coral cell type diversity and symbiosis-associated gene programmes, including in *S. pistillata*^35–38^.

Here, we generate a time- and cell-resolved transcriptomic atlas of heat stress and recovery in *S. pistillata* from the Gulf of Aqaba. We profiled coral colonies exposed to 32 °C, from early (6 h) to mid (48 h) and late stress (7 days), followed by a recovery period (45 days; Fig. 1a). This allows us to explore early and sustained cellular responses, ascertain their temporal dynamics, and quantify the extent to which different cell types and functions return to their baseline transcriptional states. Our study thus reveals how the mechanisms of stress resilience are distributed across cell types in an ecologically relevant reef-building coral.

**Figure 1.**
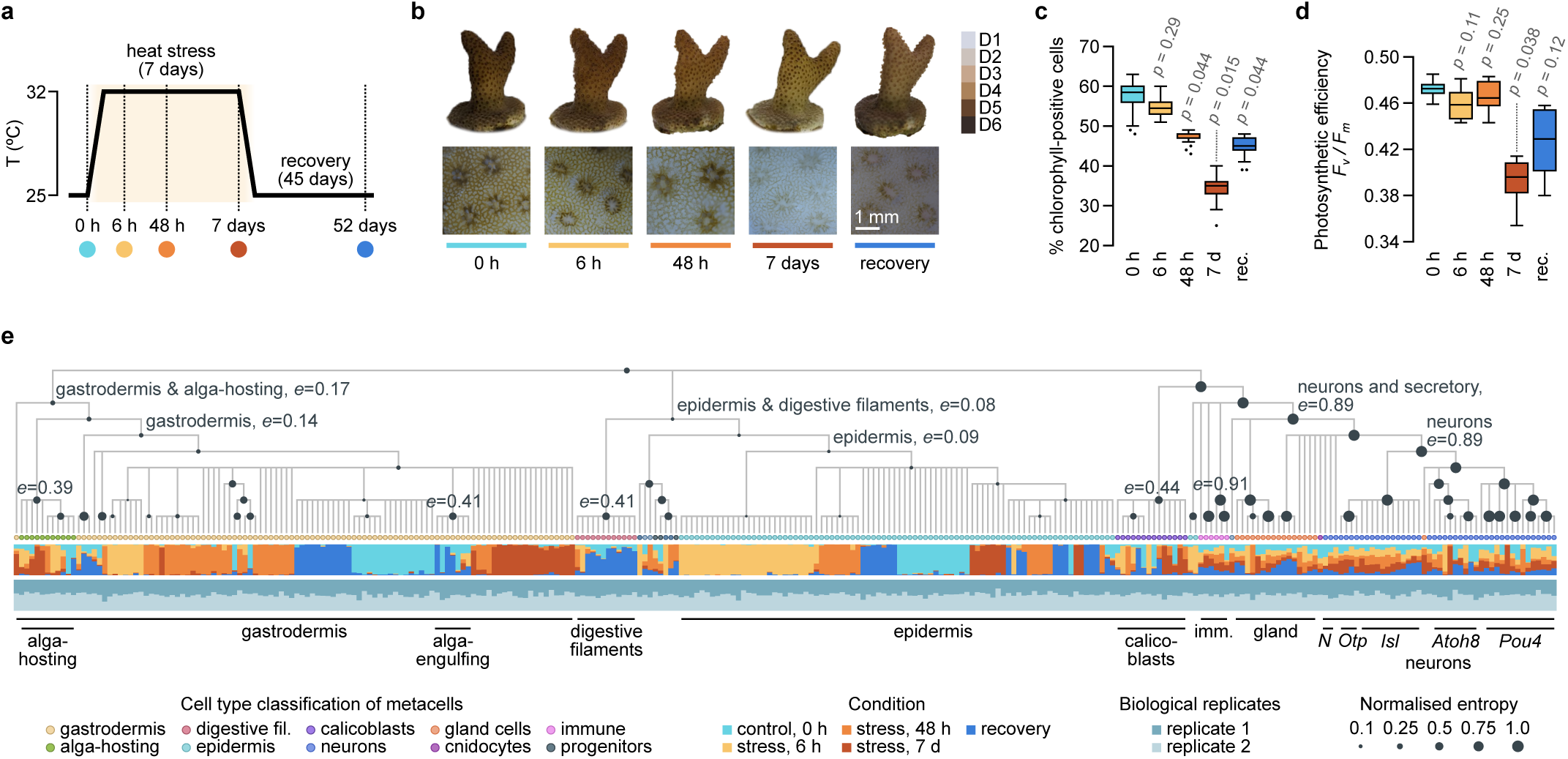
Experimental time-course of stress and recovery. **a,** Diagram of the design of the time-course of heat stress and recovery experiment, with the five sampling points used all throughout this study: control (*t =* 0 h), heat stress (*t* = 0 h, 48 h and 7 days), and recovery (after 45 days). **b,** Representative images of a *S. pistillata* nubbin at each time-point, showing the whole nubbin (top) and detailed views of the coral surface and polyps (bottom). The Coral Health Chart colorscale (*D1-D6*) is included as a visual reference for tissue colour and bleaching status. Scale bar, 1 mm. **c,** Percentage of chlorophyll-positive along the experimental time-course. Measurements were obtained from three colonies, with two nubbins per colony and three measurements per nubbin; *p-*values assess the significance of differences with *t* = 0 h measurements (*ANOVA* test, *FDR*-adjusted). **d,** Photosynthetic efficiency (*F_v_/F_m_*) along the experimental time-course, measured in the same three colonies; *p-*values assess the significance of differences with *t* = 0 h measurements (*ANOVA* test, *FDR*-adjusted). **e,** Tree hierarchical representation of transcriptional similarities between 297 metacell clusters comprising 19,830 single-cell transcriptomes (top), the time-point composition of each metacell (middle), and biological replicate composition of each metacell (bottom). Individual metacells, indicated as dots at the tree tips, are colour-coded according to their cell type identity (Extended Data Fig. 1f, g). In selected internal nodes of the tree we indicate the degree of time-point stratification of the descendant metacells, measured as a normalised entropy statistic (denoted as *e*, and by dot size; it ranges from *e =* 0 for branches where each descendant metacell is entirely composed of cells from one condition, to *e* = 1 when conditions are entirely mixed). Internal tree branches with less than 10 specific marker genes have been collapsed.

## Results

### A single-cell transcriptomic atlas of heat stress and recovery

We generated a single-cell transcriptomic time-course from ten nubbins derived from two adult *S. pistillata* colonies. Nubbins were maintained at 25 °C, and sampled at 0 h, after 6 h, 48 h and 7 days of heat exposure at 32 °C, and after 45 days of recovery at 25 °C (Fig. 1a). We imaged the specimens along the experiment (Fig. 1b), quantified their chlorophyll-positive cells (Fig. 1c), and their photosynthetic efficiency (Fig. 1d). These measurements indicate a decline in symbiont abundance and performance during heat stress, most markedly at 7 days. At this point chlorophyll-positive cells decreased from 57% to 34%, and photosynthetic efficiency from *F_v_/F_m_* = 0.47 to 0.39 relative to the control (significant at *p* < 0.04 in paired *t*-tests; Supplementary Table 1 and 2). This was followed by a partial restoration of the symbiotic state after 45 days of recovery (45% chlorophyll-positive cells, *F_v_/F_m_* = 0.43). Because the corals were maintained in a closed recirculating artificial seawater system, the restoration of the symbiont population depended primarily on the re-uptake of expelled algae, or from proliferating within host tissues. This could explain why, compared to previous experiments in open-flow seawater systems^39^, symbiont abundance and photosynthetic efficiency had not completely recovered in the final sampling point.

We profiled 19,830 single-cell transcriptomes from two biological replicates, including between 3,301 and 4,425 cells per condition (Extended Data Fig. 1a-d). To minimise batch effects, cells from different time-points were ClickTag-barcoded^40^ and pooled before single-cell profiling. We grouped the single-cell transcriptomes into 297 high-granularity clusters, termed metacells^41^, which we annotated by comparison with our previously published *S. pistillata* single-cell atlas^37^ (Extended Data Fig. 1e-g). We recovered all major cell types: gastrodermis (including an alga-hosting population), epidermis, calicoblasts (skeleton-producing epidermal cells), digestive filaments, cnidocytes (stinging cells), germline cells, and various subtypes of neurons, gland cells, and immune cells. Our biological replicates showed an even sampling of cell types (Extended Data Fig. 1h) and high concordance of the global transcriptomes at each time-point (Extended Data Fig. 1i).

Hierarchical clustering of the metacell transcriptomes broadly recapitulated cell type identity (Fig. 1e, top). Within different cell type lineages, however, metacells differed markedly in their time-point composition (Fig. 1e, middle): gastrodermal, epidermal and digestive filament metacells were strongly stratified by condition (with normalised entropy *e* ≈ 0), whereas neuronal and gland metacell clusters were highly mixed (*e* ≈ 1). Thus, the transcriptional response was strongly cell type-dependent at the metacell level: some cell populations underwent pronounced state changes during stress and recovery, whereas others remained comparatively stable. We next examined the systemic and cell type-specific gene expression programmes underlying these responses.

### Systemic responses to heat stress

To uncover systemic stress responses, we defined modules of co-expressed genes using gene-gene co-expression analysis^42^ across metacells constructed from individual time-points (47–67 metacells per condition). We identified 34 gene modules encompassing 9,696 genes (Fig. 2a), which we characterised using functional enrichment analysis (Fig. 2b, c and Supplementary Table 3). Most modules are specific to individual cell type identities (e.g. the *Atoh8+* neuron module) or groupings of related cell types (e.g. the pan-neural module, shared by multiple neuron clusters), but we also found a set of transversal modules expressed across disparate cell types (Fig. 2a). These transversal modules include basic cellular processes (e.g. protein synthesis, ciliogenesis or cell cycle progression), but also systemic responses to heat stress (Fig. 2b, c). In particular, we found five gene modules with variable expression across multiple cell types during heat stress (Fig. 2b, Extended Data Fig. 2a, b, and Supplementary Table 3).

**Figure 2.**
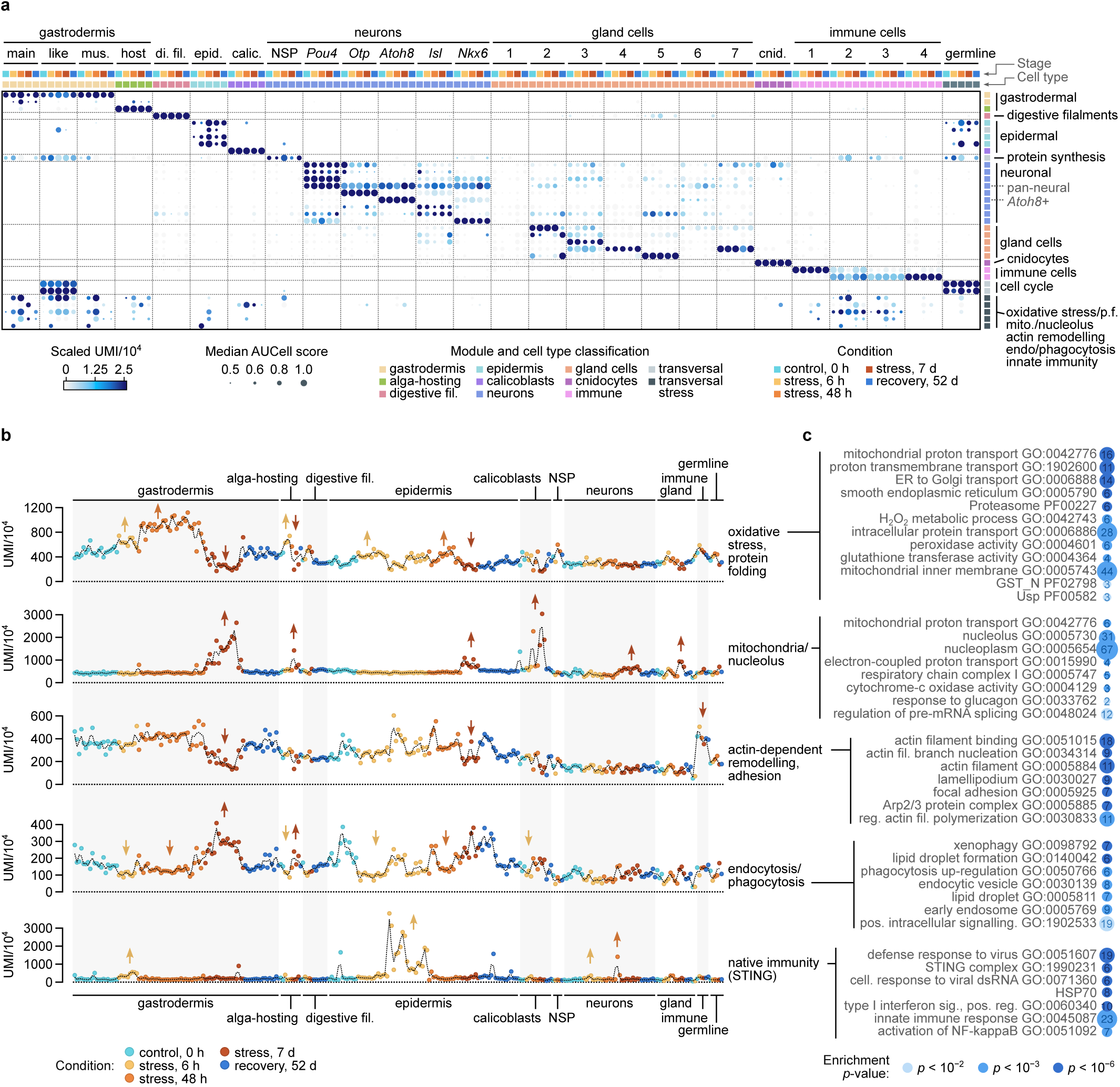
Gene module activity across cell types and conditions. **a,** Heatmap showing the activity of 34 modules of co-expressed genes (rows) in each individual cell type (columns; 26 cell types) and time-course condition (five conditions per cell type). Gene module activity in each cell population is measured as (1) the fraction of UMIs from module genes in that group of cells (scaled, colour in heatmap); and (2) its median *AUCell* score, calculated from individual cells (dot size in heatmap). Individual modules have been manually colour-coded (right side) based on their expression pattern: cell type-specific, transversal (light gray), or transversal and stress-related (dark gray). **b,** Fraction of UMIs from five transversal stress gene modules across metacell, sorted by cell type and colour-coded by time-point (from control to recovery). Dotted black lines represent a rolling average of the UMI fraction per metacell (step = 2). Coloured arrows indicate selected examples of significant up- or downregulation of a module in a specific cell type (all *p* < 2 × 10^−5^ in two-sided Wilcoxon rank-sum tests, detailed *p-*values in Supplementary Table 3). **c,** Selected enriched functional categories (Gene Ontology and PFAM domains) in the five transversal stress modules, with the number of associated genes (in-dot numbers and dot size); *p-*values are derived from one-sided hypergeometric tests for PFAM domains and Fisher’s exact tests for Gene Ontologies (colour-coded, available in Supplementary Table 3).

One broadly-expressed stress module, active in gastrodermal and epidermal cell clusters, was upregulated early during the time-course (6 – 48 h) and downregulated after prolonged exposure (7 days; Fig. 2b, c). It is enriched in H_2_O_2_ metabolism genes, mitochondrial proton transporters, peroxidases (e.g. *Prdx4*, *Prdx6*, *Sod1*, or *Sod2*), glutathione transferases, universal stress proteins (with Usp domains), genes controlling protein folding, and multiple proteasome components (Fig. 2c and Supplementary Table 3). It represents an early protective response to oxidative stress and protein misfolding.

A second transversal stress module was activated after 7 days of heat exposure in the gastrodermis, epidermis, alga-hosting cells, calicoblasts, and in various neurosecretory cell types (Fig. 2b, c). This late module includes genes associated with nucleolar function and pre-mRNA splicing, as well as additional genes involved in mitochondrial electron transport. It also includes immunity-related transcription factors (*Nfatc1-4/5* and *Nfx1*) and the circadian clock regulator *Cry1/2* (Supplementary Table 3).

At 7 days we also identified the downregulation of an actin-related module, including genes linked to the Arp2/3 complex, actin nucleation, focal adhesion, and lamellipodium formation. This was most evident in the gastrodermis, epidermis and one immune cell cluster. This downregulation could reflect a systemic decrease in actin-dependent cell adhesion, cell shape regulation and motility during prolonged heat exposure. Concurrently, gastrodermis and epidermis cells activate another late stress module with genes linked to with xenophagy and phagocytosis, early endosomes, lipid droplets, the Golgi apparatus, and intracellular signalling (Fig. 2b, c). Thus, prolonged heat stress was associated with a shift from actin-based structural and adhesion programmes towards vesicular trafficking, phagocytic and degradative pathways.

Finally, we identified an immune module activated asynchronously across cell types (e.g. 6h in the epidermis but 48 h in *Pou4+* neurons; Fig. 2b and Extended Data Fig. 2b, c). It is enriched in innate immunity genes associated with the STING pathway, NF-κB activation, response to viral double-stranded RNA, and heat-shock proteins (HSP70 domain; Fig. 2c). This module, active in both biological replicates, could reflect a viral infection linked to the host’s increased susceptibility under acute stress, or else be a fortuitous occurrence. In any event, this transcriptomic signature was transient and no longer apparent after 48 h.

Together, these gene co-expression modules reveal a set of transversal stress programmes that recur across multiple distinct cell types. Early heat exposure is marked by oxidative stress, detoxification and proteostasis responses, whereas prolonged exposure is associated with mitochondrial, nucleolar, cytoskeletal and vesicular remodelling. These systemic modules provide a common transcriptional backbone for the heat stress response, onto which cell type-specific changes are superimposed.

### Cell type-specific heat stress responses

We further characterised heat stress responses in individual cell types by differential gene expression analysis comparing each condition the control state at 25 °C. This analysis revealed substantial variability in the timing and number of differentially expressed genes (Fig. 3a). For example, the epidermis is strongly altered early in the time-course, whereas the gastrodermis, alga-hosting cells or calicoblasts showed larger transcriptional divergence after 7 days. By contrast, neuronal, gland and immune populations showed more restricted changes. After 45 days of recovery, ∼90% of the stress-responsive genes return to their baseline levels (Fig. 3b). Across all cell types, the early stress time-points (6 h and 48 h) share the largest number of activated genes, whereas the 7-day response was more distinct (Fig. 3c). A complete list of differentially expressed genes in all cell populations is available in Supplementary Tables 4 and 5 (comparing each condition to the 0 h control and to the previous time-point, respectively).

**Figure 3.**
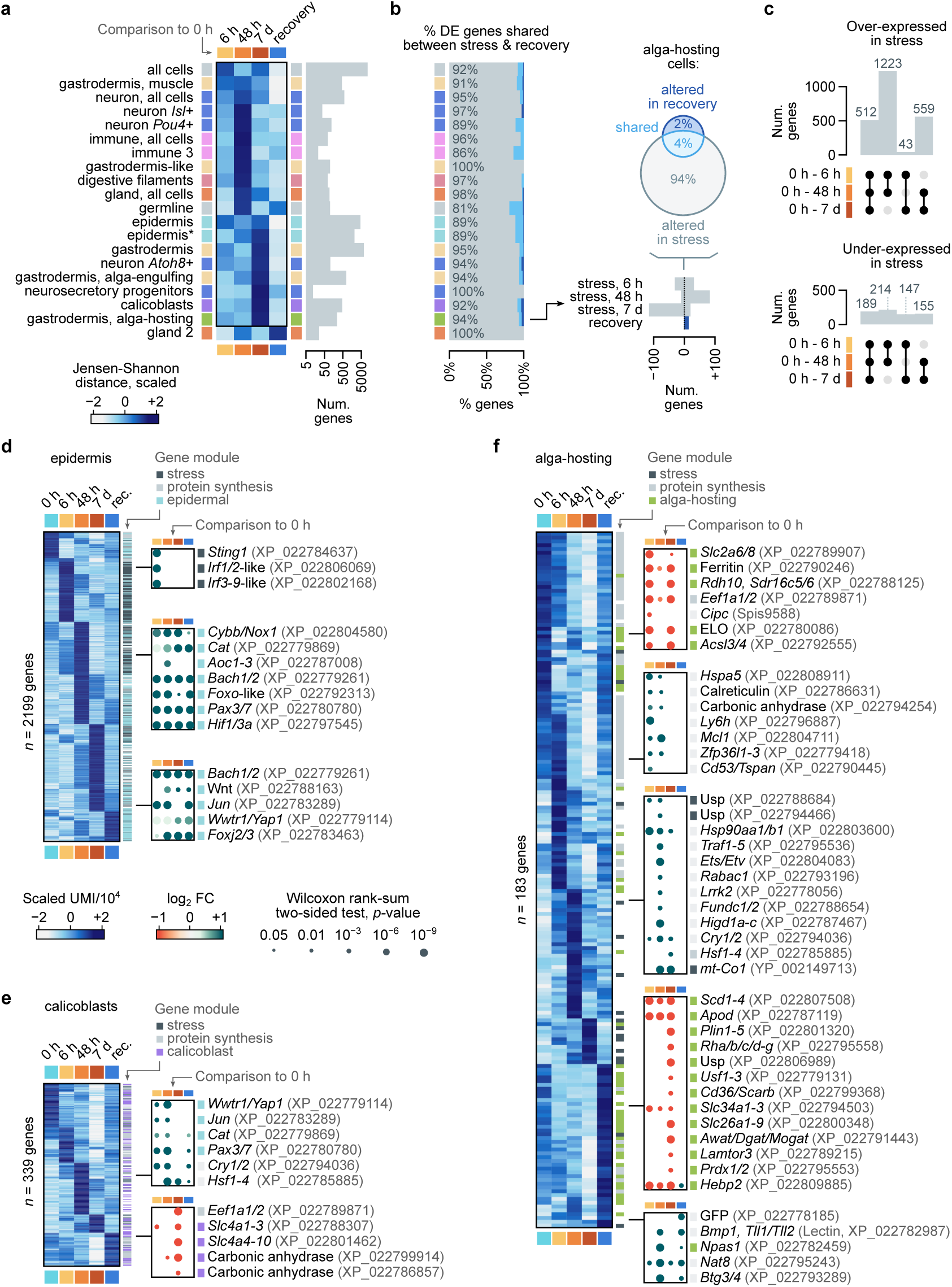
Cell type transcriptional dynamics. **a,** Scaled Jensen-Shannon distance between the transcriptome of the control condition and each later time-point (stress and recovery) in individual cell types or groups of cell types (left), and the number of differentially expressed genes considered for the Jensen-Shannon distance calculations in each cell population (right; genes included if Bonferroni-adjusted *p-*value < 0.001 and absolute fold-change > 1.1). Cell populations are sorted according to the point of maximum distance from the control (top, early; bottom, late). An asterisk indicates that genes associated with the innate immunity module have been ignored in a given cell population. **b,** Fraction of genes differentially expressed during the stress conditions (from *t* = 6 h to 7 d), the recovery condition, and both (overlap); out of all genes with altered expression in that cell population (relative to *t* = 0 h; same order as in panel *a*). The inset to the right, focusing on alga-hosting cells, shows the number of differentially expressed genes per condition (negative values indicating genes over-expressed in *t* = 0 h), and their overlap as a Venn diagram. **c,** Number of differentially expressed genes shared between two or more stress conditions (relative to *t* = 0 h), distinguishing between stress-upregulated (top) and downregulated genes (bottom). **d,** Heatmap of expression of differentially expressed genes in epidermis cells (shown as gene-scaled UMI fractions at each condition). Insets to the right indicate the fold-change (colour-coded) and statistical significance of differential expression (Bonferroni-adjusted *p-*values from Wilcoxon rank-sum tests on normalised counts; shown by dot size) for selected genes relative to the control condition. Genes are colour-coded according to selected gene module membership. **e, f,** *Id.,* for the calicoblasts and alga-hosting cell clusters.

We investigated whether the transcriptional response of each cell type was evolutionarily conserved in other corals and cnidarians, from a genetic and transcriptomic point of view (Extended Data Fig. 3a). In most cell types, the stress response was enriched in slow-evolving genes with conserved coding and *cis-*regulatory sequences, as well as conserved gene-gene co-expression profiles^43^ in other coral and anthozoan species. Coupled with the observation that stress-responsive genes in *S. pistillata* tend to be evolutionarily ancient (Extended Data Fig. 3b), this indicates that heat stress frequently mobilises evolutionarily conserved gene expression programmes. This observation also extends to the systemic stress-responsive gene modules more generally — excepting the innate immunity programme, which was enriched in young genes with fast-evolving coding and *cis*-regulatory sequences (Extended Data Fig. 3c, d).

In most cell types, the transcriptional response to heat stress is dominated by the transversal stress gene modules described above (Extended Data Fig. 3e). For example, the set of differentially expressed genes in the gastrodermis, epidermis, calicoblasts and neurons were overlapping with systemic stress gene modules (20–45% of the genes, enriched at Fisher’s exact test *p* < 2 × 10^−1^^6^): oxidative stress and protein-folding, cell motility, endocytosis/phagocytosis, innate immunity, or the mitochondrial/nucleolar module.

Nonetheless, we also detected cell type specificities in the stress responses. The epidermis showed a distinct transcriptional response dominated by the dynamic regulation of two gene modules associated with reactive oxygen species (ROS) response, and regulation of apoptosis and proliferation (Fig. 3d, Supplementary Table 3). These modules were most strongly induced after 48 h and included genes associated with ROS synthesis (e.g. *Cybb/Nox1*, or *Noxa1*), oxidative stress response (e.g. *Cat* catalases, *Aoc1-3* amine oxidases, *Noxa1,* and *Bach1/2*), apoptosis suppression (e.g induction of *Hif1/3a, Pea15a*; and downregulation of *Pidd1* and *Cradd*), tissue renewal (e.g. semaphorins, *Jun*, *Wnt1*, *Pax3/7*, *Pax1-9*-like, and *Foxj2/3*), and a *Foxo* transcription factor (a conserved regulator of stress resistance, stem cell maintenance, and longevity in cnidarians^44–46^; Fig. 3d). Between 6 and 48 h, we also observe the induction of apoptosis-suppression genes (*Hyou1*, *Hspa4*, and *Hras/Kras/Nras* GTPases; Supplementary Table 4) and the cell division gene module (Extended Data Fig. 2a). This suggests that heat stress may activate an epidermis-specific cellular state in which enhanced ROS production, potentially acting as a regulated signalling mechanism, is linked to oxidative stress regulation, tissue renewal, repair, and apoptosis avoidance.

Calicoblasts also showed a distinct, late response. After 7 days of heat stress, they strongly downregulate components of the calcium carbonate precipitation tool-kit, including carbonic anhydrases and bicarbonate SLC transporters (Fig. 3e and Extended Data Fig. 2a). This coincides with a reduced expression of protein synthesis genes and the induction of systemic stress modules, including mitochondrial and oxidative stress programmes (Fig. 3e and Extended Data Fig. 3e). This is consistent with previous studies showing reduced coral calcification and skeletal growth under elevated temperatures^47–49^, and supports a link between heat stress, disruption of the coral algal symbiosis, and impaired skeletal production.

### Stress response in alga-hosting cells

Alga-hosting cells showed one of the most distinctive responses to heat stress: their transcriptional response is not as dominated systemic stress modules, but rather by genes specific to this cell type (enriched at Fisher’s exact test *p* < 2 × 10^−1^^6^; Extended Data Fig. 3e). This response is relatively modest during the first 48 h and becomes more pronounced at 7 days (Fig. 3a), progressing from an early cell protective programme to a later loss of functions linked to the maintenance of symbiosis.

The initial response between 6 h and 48 h involved the upregulation of proteostasis genes (e.g. calreticulin chaperones, *Hspe1*, *Hspa5*, and *Ppib*) and membrane and cytoskeletal regulators (*Cd53/ Tspan*, *Ly6h* and *Dynll1*; Fig. 3f and Supplementary Table 4). This is consistent with rapid protein quality control and membrane remodelling during acute heat stress^50,51^. Alga-hosting cells also induce universal stress proteins (Usp domain), anti-apoptotic factors (*Mcl1*), and a carbonic anhydrase (involved in increasing carbon concentrations, pH regulation and inorganic carbon delivery to the symbiosome^36,52,53^). Starting at 6 h, host cells begin to repress genes associated with symbiont metabolism and nutrient storage, including glycogen synthesis (e.g. *Gys1* and *Gbe1*), lipid synthesis (ELO lipid elongases, *Acsl3/4*, and the *Srebf1* transcription factor), nutrient transporters (e.g. the sugar transporter *Slc2a6/8* or the lysosomal amino-acid transporter *Slc38a9*), intracellular trafficking (e.g. *Sort1* and *Ldlr/Lrp8/Vldlr*), and protein synthesis (e.g. *Eef1a1/2*; Fig. 3f and Supplementary Table 4).

At 48 h, the set of upregulated genes broadened markedly to include innate immunity and signalling genes (e.g. *Tirap*, *Traf1-5*, *Nlr-*family genes, *IRG*-like GTPases; Supplementary Table 4), and transcriptional regulators (*Foxo*-like, *Egr1-4*, *Deaf1*-like, *Maf*-like, and ETS transcription factors), consistent with cell state reprogramming^54,55^. Alga-hosting cells also activated genes associated with organelle trafficking and maintenance (e.g. *Rabac1, Higd1a-c*, *Fundc1*, *Chmp1b*, *Vps13d, Trappc3*, *Bloc1s2*, *Lrrk2*, *Hadha, mt-Co1* and *mt-Co2*), suggesting increased vesicle transport, membrane turnover, mitochondrial maintenance, and mitophagy-related responses.

Changes after 7 days were more profound and dominated by transcriptional downregulation (Fig. 3b, f). This included the downregulation of a carbonic anhydrase, *Rha/b/c/d-g* ammonium transporters, *Cd36/Scarb*, multiple SLC transporters, lipid metabolism genes (*Plin*, *Awat/Dgat/ Mogat*), *Prdx1/2*, and components of the Ragulator complex (*Lamtor3* and *Slc38a9*), an amino-acid sensing machinery that regulates lysosomal mTORC1 activation^56,57^. This pattern indicates loss of host transport, redox balance, lipid handling, and biosynthetic capacity, consistent with deterioration of the alga-hosting state and destabilisation of nutrient exchange ^11^. In particular, reduced inorganic phosphate and ammonium transport affects host-mediated phosphate and nitrogen availability to symbionts, which is necessary for symbiont proliferation and symbiosis stability^53,58–60^.

The persistent upregulation of the *Cry1/2* cryptochrome across heat stress (Fig. 3f), together with early repression of the circadian regulator *Cipc* (6 h) and later activation of *Npas1* (48 h), suggests that thermal stress may alter the circadian machinery of the alga-hosting cells. Because these cells normally function in close synchrony with the symbiont’s diel cycle of light-driven photosynthesis night-time metabolic reorganisation^61,62^, these changes may affect the timing of carbon exchange, nitrogen handling, and oxidative stress buffering. This temporal desynchronisation may weaken the temporal coordination required to maintain a stable symbiosis, increasing susceptibility to bleaching^63,64^.

After 45 of recovery, 94% of the heat-responsive genes in alga-hosting cells returned to their baseline state (Fig. 3b). A residual response persisted, including expression of proteostasis genes (e.g. *Dnajc25* and *Sacs*) and a green fluorescence protein-like gene (GFP). GFP-like genes have been implicated in antioxidant and photoprotective stress responses in corals, as well as in the attraction of free-living symbionts^65–67^. This host-derived cue could thus contribute to tissue protection and re-establishment of symbiosis after bleaching. In parallel, *Hebp2* was lowly expressed throughout heat stress but upregulated in recovery, consistent with the restoration of a mitochondrial apoptotic regulator. This transient repression likely functions as a host pro-survival mechanism to delay cell death and preserve tissue integrity, aligning with established cnidarian-specific strategies where apoptotic pathways are strictly modulated to dictate the tipping point between survival and bleaching^68,69^.

Together, these data indicate that alga-hosting cells follow a staged response to heat stress. During acute exposure, they activate chaperone, anti-apoptotic and membrane-remodelling programmes while retaining parts of the symbiosis-related host-cell state. With prolonged stress, this response shifts towards repression of transport, lysosomal, lipid-processing and carbon- and ni-trogen-exchange genes that are characteristically alga-hosting^1,11,36,37^. Thus, bleaching is preceded by a progressive erosion of the transcriptional programme that supports intracellular symbiosis, followed by an incomplete but substantial restoration of the pre-stress transcriptional state after recovery.

### Transitional alga-hosting cell states

The whole-organism time-course highlights alga-hosting cells as one of the most stress-disturbed cell populations. To resolve their transcriptional heterogeneity at higher resolution, we performed a targeted analysis of alga-positive cells, isolated by fluorescence-activated cell sorting (FACS; Fig. 4a, b) and sequenced using MARS-seq along the same experimental time-course used for the whole-organism single-cell atlas. We retained for further analyses the single-cell transcriptomes containing signal from both host cells and their symbiont, *Symbiodinium microadriaticum* (Extended Data Fig. 4a-c).

**Figure 4.**
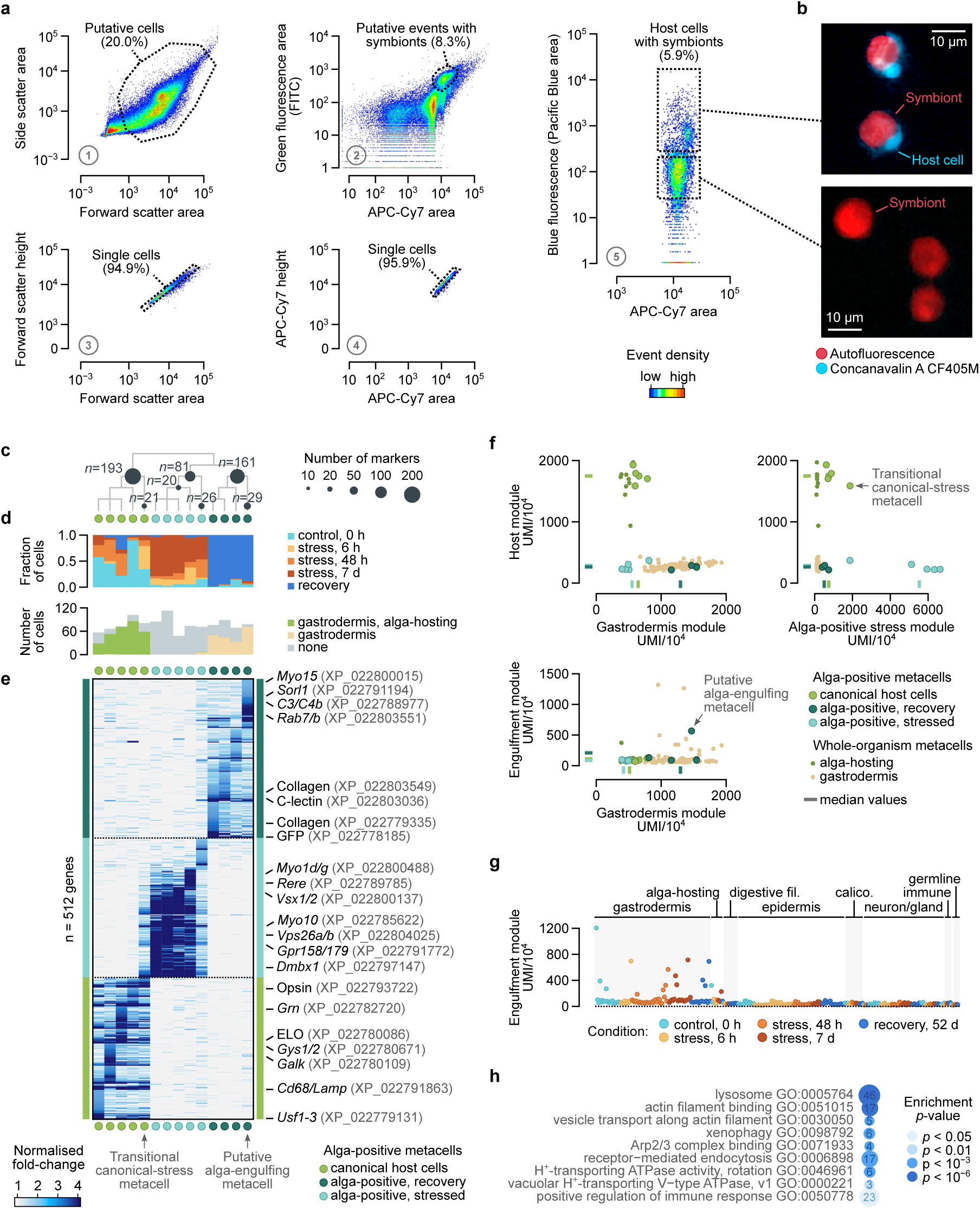
Heterogeneity in alga-hosting cells. **a,** Representative gating strategy used to isolate single coral host cells containing intracellular algal symbionts. Initial gates were applied to select putative cellular events based on forward scatter area and side scatter area (step 1), followed by exclusion of non-cellular particles by retaining DRAQ5-positive events (step 2). Doublets and multiplets were removed by comparing forward scatter area and height (step 3), and APC-Cy7 area and height (step 4). Host cells containing algal symbionts were then enriched by selecting single DRAQ5-positive events with Concanavalin A CF405M blue fluorescence, marking coral cell membranes, together with APC Cy7 autofluorescence, corresponding to chlorophyll autofluorescence from Symbiodiniaceae (step 5). **b,** Representative fluorescence images of sorted events, showing host cells with symbionts (top) and free symbionts (bottom). Coral host cells are labelled by Concanavalin A CF405M blue fluorescence, while Symbiodiniaceae symbionts are identified by red chlorophyll autofluorescence. Scale bar, 10 µm. **c,** Hierarchical tree representation metacell transcriptomes from alga-positive cells (14 metacells in total). We indicate the number of node-specific marker genes in internal nodes of the tree (*n*; denoted by dot size), which delineate three main cell populations (colour-coded dots at the tree tips). **d,** Time-course condition composition of each metacell (top) and number of cells per metacell assigned to reference cell types (assignment probability ≥ 95%) or none (< 95%; bottom). **e,** Heatmap of expression of 258 genes in each alga-positive metacell (showing up to 40 genes per metacell with normalised fold-change ≥ 2), with selected markers highlighted to the right. **f,** Fraction of UMIs in each alga-positive metacell from genes assigned to the modules characteristic of alga-hosting cells, gastrodermal cells, and the engulfment gene module (gene modules defined in the whole-organism metacell set; Fig. 2a), and the alga-positive stressed cells (defined from the alga-positive metacell set). Large coloured dots denote metacells from the alga-positive set (with median values as thick lines); smaller dots denote metacells from the gastrodermis re-clustering dataset. **g,** Fraction of UMIs from the engulfment gene modules across metacells, sorted and colour-coded by cell type (non-gastrodermal cells in gray). **h,** Selected Gene Ontology terms enriched in the genes differentially over-expressed (Bonferroni-adjusted *p-*value < 0.001 and fold-change > 1.1) in metacells with active engulfment gene module (standardised module eigenvalue *Z* > 1.28). For each Gene Ontology term we indicate the number of associated genes (in-dot numbers and dot size); *p-*values are derived from Fisher’s exact tests for Gene Ontologies (colour-coded, available in Supplementary Table 6).

We analysed 1,082 validated alga-positive cell transcriptomes, which we clustered into 14 metacells that could be grouped into three populations defined by hierarchical clustering and specific marker genes (Fig. 4c). As expected, one population showed strong expression of the canonical alga-hosting programme, including genes associated with lipid metabolism, galactose metabolism and host-symbiont nutrient exchange (Fig. 4d-f). These cells are mostly associated with the host cell cluster identified in the whole-organism atlas (Fig. 4d).

We also identified a second population of alga-positive cells, mostly from late stress (7 days), that did not map confidently to any cell type in the whole-organism atlas (Fig. 4c, d). These cells lacked expression of the canonical alga-hosting gene programme but showed a concerted expression of a distinct gene module (Fig. 4f). Their markers include specific transcription factors (a *Dmbx1* paralog, *Vsx1/2* and a *Rere* cofactor; Fig. 4e) and genes involved in endosomal transport and recycling (e.g. *Sorl1*, *Lyst*, *Vps26a/b*, *Dennd5a/b*, *Mcoln1/2/3*, *Nsf*, *Inpp5f*, and *Osbpl1/2*), actin-mediated vesicle transport (e.g. unconventional myosins *Myo10* and *Myo1d/g*), and light-sensing proteins (e.g. *Vsx1/2*, *Bco/Rpe65,* and two GPCRs, G*rm1-8* and *Gpr158/179*). We interpret this rare 7-day population as a stress-linked transcriptional state in which formerly nutrient-exchanging host cells are recycling symbiosome components. Consistent with this interpretation, we identified a transitional metacell enriched at 6h and showing intermediate expression of both the canonical and stressed modules (Fig. 4e, f).

Lastly, a third group of alga-positive cells was enriched in the recovery condition (Fig. 4c, d) and showed high expression of the general gastrodermal gene module (Fig. 4f), including key marker genes of the post-stress gastrodermis such as collagens, ependymins, and GFP (Fig. 4e and Supplementary Table 6). These cells express multiple genes associated with extracellular-matrix remodelling and tissue re-patterning, suggesting a coordinated Wnt, BMP, and TGF-β-like programme (e.g. *Wnt*, *Wif1*, *Bmper*, *Smad6/7* and *Tgfbr3*; transcriptional regulators *Egr1-4*, *Ets1/2*, *Klf5*, and *Maf*; and remodelling signalling genes such as *Agrn, Cilp*, *Cthrc1*, *Timp1*, and *Cd248*; Fig. 4e and Supplementary Table 6). These could be interpreted as matrix-remodelling gastrodermal cells involved in regenerative or tissue-patterning processes that emerge specifically during recovery from heat stress^70–72^.

This analysis also highlighted a rare metacell (F,ig. 4e and Extended Data Fig. 4e) expressing a distinctive gene module that, in the whole-organism atlas, was active only in a small sub-population of gastrodermis metacells (Fig. 4f, g). A differential expression analysis against the non-host gastrodermis indicated that this cell state is consistent with the early stages of symbiosome formation or new symbiont engulfment (Fig. 4h, Extended Data Fig. 4f, and Supplementary Table 6). For example, they showed high expression of many lysosomal sorting genes (e.g. *Cd68/Lamp* and *Sort1*), V-ATPase proton pumps that create the acidic environment of the symbiosome, and many vesicle-trafficking genes such as myosins (*Myo15*, *Myo1c/h*, *Myo1e/f*, *Myo5a-c*, and *Myo10*), Rab GTPases (*Rab7/b*), immune modulators (e.g. *Ifi35*, *Ifi44*, *Cd27/Cd40*, *Rsad2* and *Znfx1*), and microbial recognition genes (*Bpi*, *Lbp* and *Mpeg1*). They also express a number of proteins linked to host-symbiont cell interactions in other corals^73,74^, such as complement genes (including *C3/C4b*), scavenger receptors, and secreted C-type lectins (Extended Data Fig. 4f). Interestingly, this includes multiple paralogs of *LePin* (Extended Data Fig. 4g), a secreted lectin from the octocoral *Xenia* sp. that binds to and promotes the phagocytic uptake of algae during the early establishment of symbiosis ^35,75^. By contrast, these cells lacked the transcriptional programme associated with a stable metabolic integration of the symbiont (Fig. 4f). This profile is consistent with proximal host-symbiont cell interactions, phagocytic uptake of symbionts, symbiosome formation and maturation, and the continued coupling of intracellular trafficking with innate immune regulation during recovery, as described for coral and other cnidarian symbioses^1,36,74^.

## Discussion

Our single-cell time course reveals coral heat stress as a temporal progression from early cellular protection to late tissue and symbiosis remodelling, followed by partial recovery. During early phases of heat exposure, the dominant response is protective and broadly conserved: multiple cell types activate oxidative-stress, detoxification and proteostasis programmes, including chaperones, antioxidant enzymes and protein-quality-control pathways (Fig. 2b, c). This systemic response is accompanied by early cell type-specific programmes: epidermal cells activate ROS-associated, tissue-renewal and apoptosis-avoidance genes, whereas alga-hosting cells induce proteostasis, survival and membrane-regulatory programmes while beginning to repress genes involved in nutrient storage, lipid metabolism and transport (Fig. 3). Thus, acute heat stress does not immediately erase host cellular identity. Instead, it initiates a coordinated stress-buffering response while beginning to weaken the metabolic programme that supports intracellular symbiosis.

After 7 days of heat exposure, the response shifts from protection to cellular reorganisation and symbiosis destabilisation. Across multiple tissues, prolonged stress is associated with mitochondrial, nucleolar and RNA-processing programmes, reduced actin-dependent adhesion and cell-shape regulation, and activation of vesicular trafficking, phagocytic and degradative pathways (Fig. 2b, c). These late systemic responses coincide with pronounced tissue-specific effects. Calicoblasts repress carbonic anhydrases and bicarbonate transporters linked to calcium carbonate precipitation, providing a cellular link between heat stress and impaired skeletal formation (Fig. 3e). In alga-hosting cells, prolonged stress represses major components of the host symbiotic programme, including nutrient exchange, lipid storage and processing, redox control, lysosomal nutrient sensing and intracellular trafficking (Fig. 3f). This indicates a progressive loss of the cellular functions that maintain metabolic integration with intracellular symbionts. Thus, prolonged heat stress moves the coral from homeostatic buffering towards active dismantling of the cellular machinery that sustains calcification and intracellular symbiosis. Importantly, the gene expression programmes mobilised during heat stress are enriched in evolutionarily ancient, well-conserved genes, both genomically and transcriptomically (Extended Data Fig. 3a-c).

Coral recovery involves a substantial reversal of the stress-induced transcriptomic changes, with ∼90% of genes returning to baseline levels after 45 days at 25 °C (Fig. 3a, b). The residual post-stress signature is marked by persistent proteostasis, chaperone-related quality control, RNA processing adjustment, and limited metabolic remodelling. But we also observe an incomplete functional restoration of the symbiotic partnership, as algal abundance and photosynthetic efficiency recover only partially (Fig. 1b, c). This suggests a possible temporal uncoupling in the recovery process of the host and symbiont cells, particularly when access to new free-living algae is limited. The recovery stage also contains alga-positive gastrodermal states associated with extra-cellular-matrix remodelling, Wnt, BMP and TGF-β-like signalling, and a rare state consistent with symbiont engulfment and early symbiosome formation.

These findings frame bleaching and recovery as dynamic cellular transitions. Conserved stress programmes preserve host viability, late stress programmes dismantle key coral functions (symbiosis and calcification), and recovery programmes rebuild tissue architecture and host-symbiont interactions. By resolving these processes at cellular resolution, this atlas provides a reference for identifying the host states and pathways that shape bleaching outcomes. Future work should test whether the stress and recovery states described here are conserved across coral species with different thermal sensitivities, across symbiont lineages and under combined environmental stressors, including heat and acidification. With global marine heat waves increasing in severity and affecting an ever wider range of corals^17,23,24^, comparative analyses of the heat stress responses in multiple species will become a key tool to understand their adaptation mechanisms. Furthermore, coupling single-cell profiling with spatial transcriptomics, live imaging, physiological measurements and targeted perturbations will be needed to connect these transcriptional states to symbiont loss and reacquisition, symbiosome recycling, and calcification. These comparative and functional approaches may help explain why some coral holobionts recover whereas others fail, and provide a foundation for strategies aimed at preserving coral reef function under increasingly frequent marine heat waves.

## Methods

### Coral collection, maintenance, and experimental design

Three colonies of *Stylophora pistillata* were collected from the Gulf of Aqaba, offshore from the Interuniversity Institute for Marine Sciences in Eilat, Israel (collection permit issued by the Israel Nature and Parks Protection Authority #2022/42970). Colonies were acclimated for two weeks and subsequently fragmented into nubbins. The nubbins were allowed to recover and acclimate for an additional four weeks before the onset of the heat stress experiment. The experiment was conducted in a controlled aquarium system at the Leon H. Charney School of Marine Sciences, University of Haifa. Corals were maintained in artificial seawater prepared with Red Sea Salt, Red Sea Ltd, at a salinity of 39 ppt and 25 °C, under a 12 h:12 h light:dark photo-period at a photosynthetically active radiation level of 50 μmol photons m^−2^s^−1^. Corals were fed twice per week with planktonic coral food (Reef Snow, Brightwell Aquatics), according to the manufacturer’s instructions.

The heat stress experiment was performed using three aquaria. Coral nubbins derived from the three parental colonies were randomly distributed among the aquaria before the start of the experiment. The experiment included five sampling time-points: baseline control at 25 °C, designated 0 h, followed by 6 h, 48 h and 7 days of heat exposure at 32 °C, and a final recovery time-point after 45 days at 25 °C. At the onset of the experiment, water temperature increased from 25°C to 32 °C over 3 h. The 6 h time-point was collected 6 h after the start of the temperature ramp, corresponding to 3h after reaching 32 °C. Corals were maintained at 32 °C for 7 days, after which the temperature was returned to 25 °C for a 45 days recovery period. The recovery samples were therefore collected 52 days after the beginning of the experiment. At each point, coral nubbins were sampled for cell counting, photosynthetic efficiency measurements, and single-cell transcriptomic profiling.

At each experimental point, nubbins were photographed together with a Coral Health Chart colour reference card (*D* scale)^76^ to provide a standardised visual record of colony colour and to monitor bleaching progression throughout the heat stress and recovery experiment. For quantification of bleaching, two nubbins from each of the three independent parental colonies were collected at each time-point. These samples were used to quantify total coral host cell abundance and Symbiodiniaceae cell abundance. Each fragment was measured using three repeated counts. Photosynthetic efficiency was monitored throughout the experiment at all five time-points using two nubbins from each of the three independent parental colonies (see below). The same sampling structure was maintained across time-points, with two repeated measurements collected per fragment. For single-cell transcriptomic analysis (see below), two nubbins were collected at each time-point from each of two parental colonies. The same two colonies were used for single-cell transcriptomic profiling across all sampled time-points.

#### Quantification of Symbiodiniaceae cell abundance

Symbiont abundance was quantified at each experimental time-point using two nubbins from each of the three independent parental colonies, yielding six nubbins per time-point. Nubbins were processed individually. Each nubbin, approximately 1–2 cm in length, was first washed three times in filtered 0.22 μm calcium free artificial seawater, CaF ASW, containing 10 mM Tris HCl, pH 8, 2.1427 mM NaHCO_3_, 10.7309 mM KCl, 426.0123 mM NaCl and 7.0403 mM Na2SO4. After washing, each nubbin was transferred to a sterile well of a 6 well plate containing 10 ml CaF ASW supplemented with 0.5 mM EDTA and incubated for 10 minutes at room temperature, following the tissue dissociation conditions previously described for *S. pistillata* single-cell preparation^36^. Coral tissue was mechanically dissociated from the skeleton by scraping the nubbin surface with a sterile 10 μl plastic pipette tip, allowing removal of tissue from within the polyp cavities. Each nubbin and its cell containing CaF ASW solution were then transferred to a 50 ml tube, with sufficient solution to fully cover the nubbin. Tubes were placed on an orbital shaker for an additional 20 minutes at room temperature, with vigorous pipetting every few minutes, until the skeleton was visually clean of coral tissue. The skeleton was then removed, and the resulting cell suspension was used for cell counting. Cell suspensions were counted using an Invitrogen Countess II FL Automated Cell Counter. Total cell abundance was measured in brightfield mode, while algal symbionts were detected using chlorophyll autofluorescence in the fluorescence channel. For each nubbin, three repeated counts were acquired, and the percentage of Symbiodiniaceae was calculated as the number of chlorophyll autofluorescent cells divided by the total number of cells counted in brightfield.

The statistical significance of symbiont abundance variation across conditions was assessed using ANOVA tests, with each coral specimen as an independent replicate, and pairwise Welch two-sample *t*-tests, correcting for paired comparisons using the Benjamini-Hochberg *FDR* procedure (Supplementary Table 1).

#### Photosynthetic efficiency measurements

Photosynthetic efficiency was assessed at each experimental time-point using a Diving Fluorescence Induction and Relaxation (FIRe) fluorometer^77,78^. For these measurements, two dedicated nubbins from each of the three independent parental colonies were used, yielding six measured nubbins per time-point. These nubbins were reserved specifically for fluorescence measurements and were not used for single-cell transcriptomic profiling, to prevent any potential effect of the measurement light exposure on transcriptional responses. Prior to measurement, nubbins were dark acclimated for 20 minutes. Each nubbin was measured twice at randomly selected positions on the coral surface. The FIRe system records chlorophyll fluorescence induction and relaxation profiles and derives the maximum quantum efficiency of photosystem II, *F_v_/F_m_*, from the minimum and maximum fluorescence yields, *F_o_* and *F_m_*. *F_v_/F_m_* was used as an indicator of photosynthetic performance of the coral associated Symbiodiniaceae during heat stress and recovery^79^.

The statistical significance of changes in *F_v_/F_m_* across conditions was assessed using ANOVA tests, and pairwise Welch two-sample *t*-tests, correcting for multiple comparisons with the Ben-jamini-Hochberg *FDR* procedure (Supplementary Table 2).

### Single-cell transcriptomics

#### Specimen dissociation and cell fixation

To dissociate and fix coral cells for 10x scRNA-seq, we used a modified version of the ACME maceration protocol^80^. For scRNA-seq, cells were dissociated from nubbins collected from two colonies, representing two biological replicates, across all five experimental conditions. In total, 10 nubbins were processed, with one nubbin per colony for each condition. Coral nubbins, 2–3 cm in length, were washed with filtered (0.22 μm) CaF-ASW (10 mM Tris-HCl pH 8, 2.1427 mM NaHCO3, 10.7309 mM KCl, 426.0123 mM NaCl, 7.0403 mM Na2SO4) and transferred to a 50 ml tube containing 10 ml of ACME maceration solution without BSA, ensuring that the solution fully covered the nubbins. The ACME solution was prepared as follows: 6 ml CaF-ASW, 1 ml glacial acetic acid, 1 ml glycerol, 1.5 ml methanol, and 0.5 ml EDTA (a 13:2:2:3:1 ratio). Samples were incubated at room temperature for 30 minutes with periodic pipetting. The cell suspension was then filtered through a 70 μm strainer into a new 50 ml tube, kept on ice, and aliquoted into 1.5 ml portions in 2 ml tubes. Aliquots were centrifuged at 1,000 g for 10 minutes at 4°C. The supernatant was discarded, and the pellet was resuspended in 1 ml of Resuspension Buffer 1 (RB1), prepared by mixing 1 ml 10× PBS, 3.3 ml 2.4 M sorbitol, 5.7 ml nuclease-free water, and 20 μl RNAsin. Cells were washed again with RB1 buffer under the same centrifugation conditions. Cells were counted using DAPI staining (9 μl of cell suspension mixed with 1 μl DAPI, 1 mg/ml). Cell concentration was calculated by multiplying the average cell count from four sets of 16 squares by 10,000. The sample volume was adjusted to obtain aliquots of 100 μl containing 400,000 cells each.

#### Clicktag barcoding for 10x scRNA-seq

Fixed cells were barcoded using a modified version of ClickTags^40,81^. To optimise the labelling reaction in ACME fixative, the amine reactive cross-linker TCO-NHS used by Gehring *et al.*^40^ was replaced with TCO-PEG4-TFP (Click Chemistry Tools), which has improved stability against hydrolysis in aqueous media. Barcoding DNA oligonucleotides (ClickTags) carrying a 5′ amino modifier (Integrated DNA Technologies) were activated by derivatisation with methyltetrazine NHS-ester, as originally described^40^. For cell tagging, each time-point sample was labelled using a unique combination of three MTZ-derivatised oligonucleotides. Cell suspensions were pre-incubated with 4.5 μl of 1 mM TCO-PEG4-TFP for 5 minutes at room temperature, protected from light. Premixed MTZ-activated tags (12 μl total) were then added, followed by thorough mixing. Samples were incubated for 30 minutes at room temperature on a rotating platform, protected from light. The reaction was quenched by adding 13 μl of 100 mM Tris-HCl, resulting in a final concentration of 10 mM, and 0.65 μl of 10 mM MTZ-DBCO. Samples were incubated for an additional 5 minutes at room temperature. Each pool was mixed with two volumes of RB1, inverted three times and centrifuged at 1,000 g for 10 minutes at 4°C. The pellet was washed with 1 ml RB1 and centrifuged again under the same conditions. Finally, cells were resuspended in 900 μl RB1 and 100 μl DMSO. Samples were stored at −80 °C until sorting for scRNA-seq.

#### Cell sorting and 10x scRNA-seq

Single-cell transcriptomes were generated using the Chromium Single Cell n3′ Gene Expression kit v3.1, 10x Genomics. Frozen ClickTag barcoded samples were thawed on ice, transferred to 1.5 ml tubes and washed in RB2, containing 1× PBS, 0.5% BSA and RNase inhibitor. Cells were pelleted by centrifugation at 1,000–1,500 g at 4°C, washed again in RB2 and resuspended in 1 ml RB2 before sorting. Cells were stained with DRAQ5 (1:330; Thermo Fisher Scientific, 62251) to label nuclei, and Concanavalin A CF405M (1:400; Biotium, 29074), to label coral cell membranes. Cells were sorted using a FACSAria II SORP cell sorter equipped with a 70 μm nozzle and operated in single-cell purity mode. Non-cellular particles were excluded by selecting DRAQ5-positive events, and doublets and multiplets were removed using forward scatter parameters. To enrich for coral host cells containing algal symbionts, cells were selected based on DRAQ5 positivity, C1oncanavalin A CF405M signal and Cy7 autofluorescence, corresponding to chlorophyll autofluorescence from the algal symbionts (Fig. 4a, b). Sorting gates were calibrated using dedicated control samples and gates established from MARS-seq sorting plates. For each 10x reaction, five time-point samples were pooled by FACS sorting into a single well containing the 10x reverse transcription reagent mix. A total of 40,000 events were sorted per reaction, corresponding to 8,000 events from each time-point sample, with the aim of recovering approximately 15,000 cells after 10x loading. After sorting, RT Enzyme C was added, the sample was mixed by pipetting, adjusted to the required loading volume with nuclease-free water when necessary, transferred to an ice-cold Protein LoBind tube, and immediately encapsulated using the 10x Chromium controller. Barcoded cDNA and gene expression libraries were prepared according to the 10x Genomics protocol. For ClickTag library preparation, ClickTag cDNA was recovered from the supernatant of the first 0.6× SPRI purification after cDNA amplification. The recovered ClickTag cDNA was treated with Exonuclease I, purified with AMPure beads and amplified by PCR using up to 7 ng of input cDNA. ClickTag libraries were purified by double size selection using SPRI beads, and final library size distribution and concentration were assessed using TapeStation and Qubit.

#### Cell sorting and Massively Parallel Single-Cell RNA-seq (MARS-seq)

Cells were dissociated and fixed as described above (except for the ClickTag barcoding steps) and stored at −80 °C until sorting for MARS-seq. Frozen samples were thawed on ice, and cells were collected by centrifugation at 2,000 g for 5 minutes at 4°C. After washing with 2 ml RB2, cells were pelleted again and resuspended in 3 ml RB2. Cells were stained with DRAQ5, 1:330 (Thermo Fisher Scientific, 62251) to label nuclei, and Concanavalin A CF405M, 1:400 (Biotium, 29074) to label coral cell membranes. Cells were sorted using a FACSAria II SORP cell sorter and distributed into 384-well capture plates from the same plate production batch, with each well containing 2 μl lysis solution composed of 0.2% Triton X-100, RNase inhibitors and barcoded poly(T) reverse transcription primers for scRNA seq. For MARS-seq, only coral host cells containing algal symbionts were targeted. Non-cellular particles were excluded by selecting DRAQ5-positive events, and doublets and multiplets were removed using forward scatter width (FSC-W) versus forward scatter height (FSC-H). From the singlet population, cells positive for DRAQ5, Concanavalin A CF405M and Cy7 were sorted, with the Cy7 signal corresponding to chlorophyll autofluorescence from the algal symbionts. For each time-point, three complete 384-well plates were sorted, corresponding to 1,152 host cells containing Symbiodiniaceae. Sorted plates were immediately centrifuged at 800 g to ensure immersion of cells in the lysis solution, placed on dry ice and stored at −80 °C until further processing. Libraries were prepared in parallel using MARS-seq, as previously described^82^. Briefly, mRNA was converted into cDNA on a Bravo automated liquid handling platform, Agilent, using oligonucleotides containing both unique molecular identifiers and cell barcodes. PEG8000 was added to the reverse transcription reaction at a final concentration of 0.15% to increase cDNA capture efficiency, and unused oligonucleotides were removed by exonuclease I treatment. cDNA was pooled, with each pool representing one 384-well MARS-seq plate, and linearly amplified by T7 in vitro transcription. The resulting RNA was fragmented and ligated to an oligonucleotide containing the pool barcode and Illumina adaptor sequences using T4 ssDNA:RNA ligase. RNA was then reverse transcribed into DNA and PCR amplified. The size distribution and concentration of the resulting libraries were assessed using TapeStation (Agilent), and Qubit (Invitrogen). scRNA seq libraries were pooled at equimolar concentration and sequenced to saturation (≥ 6 reads/UMI) on an Illumina NextSeq 500, using high output 75 cycle v2.5 kit (Illumina). In total, we obtained 190 million reads, with a median of 22,000 uniquely mapped reads per cell.

#### Mapping of whole-organism single-cell 10x transcriptomic dataset

Both 10x cDNA libraries were mapped to the reference genome with *CellRanger*^83^ 7.2.0 to retrieve matrices of counts of unique molecular identifiers (UMI) per gene and cell. We used whole gene regions to count UMIs in each genome (*--transcriptome* parameter), which were extended to include proximal downstream regions so as to compensate for the low-quality annotation of UTRs in this species^36,84^. We have used the same 3′ gene extensions used in our previous studies on *S. pistillata*^36,37^.

#### Filtering of single-cell transcriptomes by removal of doublet and empty droplets

We used the distributions of unique molecular identifies (UMI) per cell in each of the two 10x single-cell scRNA-seq experiments to identify *bona fide* cells by visual inspection and selection of a threshold to distinguish cells from empty droplets (≥ 300 UMI in both cases; Extended Data Fig. 1b).

We also used Clicktag-multiplexed libraries to experimentally identify doublet cells in the two 10x single-cell scRNA-seq experiments, as described previously^37,81,85^. Specifically, we sequenced ClickTag libraries from the same cells used in the 10x experiments and used ClickTag barcode counts to identify doublet cells in the overloaded cDNA experiments by assigning three unique barcodes to each sample, as follows: (i) mapped the Clicktag-derived read set to an *in silico* synthetic transcriptome consisting of all barcode sequences with all possible site-wise substitutions, using *kallisto*^86^ 0.46.2 for mapping (*bus* module), correction, sorting and counting (*correct*, *sort*, and *count* modules); (ii) we retained cells with at least 10 total counts in the top-scoring barcode combination; (iii) we normalised the barcode counts (divided by the total number of counts of each barcode) and compared the ratio of the sum of normalised counts from each set of same-sample barcodes to the second most abundant set, retaining cells with a first-to-second ratio >2; (iv) recorded the three most abundant barcodes in each cell and retained the cells for which all of them were associated with the same initial sample (Extended Data Fig. 1c). Cells with sufficient counts, concordant top barcodes, and first-to-second normalised count ratios > 2 were classified as singlets, assigned to each of the five stress time-points, and retained for downstream analysis (Extended Data Fig. 1d). Cells with insufficient ClickTag counts or sufficient counts but discordant top barcodes were flagged as unclassifiable or doublets, respectively, and removed from downstream analyses.

We further removed cells with very low number of unique genes. Specifically we discarded cells with low number of unique genes in a scaled distribution of log-transformed counts per cell (*z*-score < −3.09, i.e. *p* < 0.001; this resulted in the elimination of 32 cells with less than 104 genes).

#### Clustering and annotation of single-cell transcriptomes

We concatenated the two filtered UMI matrices from each 10x experiment (i.e. after removing doublets or non-cell droplets, see above) in a single *Seurat R* object^87^ (version 5.3.0), retaining for each cell the experiment of origin (as batch information) and its stress condition, from 0 h to 52 days (as inferred from the combination of top ClickTag barcodes in the pooled sequencing experiments). The UMI counts matrix was normalised using the *Seurat::NormalizeData* function (parameters: *normalization.method = “LogNormalize”*, *scale.factor = 1e4*).

In order to annotate the cell type of origin of each cell, we integrated the heat stress dataset with our previously published *S. pistillata* 10x single-cell transcriptomic atlas^37^. First, we concatenated the UMI matrices of both studies and constructed high-granularity metacell clusters on the joint atlas using the *metacell*^41^ 0.3.7 *R* package, as follows: (i) we calculated gene variability statistics with *metacell::mcell_add_gene_stat*, (ii) selected variable features with *metacell::mcell_gset_filter_multi* (parameters: *T_tot = 100, T_top3 = 2, T_szcor = −0.05, T_niche = 0.01*, and blacklisting all ribosomal genes with *blacklist*, defined from Pfam domain content); *(iii*) calculated the graph of best cell-cell neighbours with *metacell::mcell_add_cgraph_from_mat_bknn* (parameter *K = 100*); (iv) calculated the co-clustering matrix from resampling the graph, *with metacell::mcell_coclust_from_graph_resamp* (using a minimum metacell size of *min_mc_size = 10*, and 100 re-samplings of 75% of the genes with *p_resamp = 0.75* and *n_resamp = 100*); (v) constructed metacell clusters from the balanced co-clustering profiles with *metacell::mcell_mc_from_coclust_bal-anced* (*K = 30* cells, minimum metacell size of *min_mc_size = 10* cells, *alpha* parameter = 2).

In parallel, we built low-granularity Leiden clusters with the *Seurat::FindClusters* function (parameters: *resolution = 4, algorithm = 4, method = “igraph”*) applied to a *Harmony*^88^-corrected (*Seurat::In-tegrateLayers* with parameters *method = HarmonyIntegration, normalization.method = “SCT”* and *k.-weight = 50*) dimensionality reduction (principal component analysis with *Seurat::RunPCA*with *npcs = 100*, only the top 19 PCs were retained based on the first derivative criterion applied to the fraction of explained variance) of the SCT-transformed^89^ count matrix (i.e. variance stabilising transformation with *Seurat::SCTransform* function, with *do.scale = TRUE* and *do.center = TRUE*).

This resulted in 546 preliminary metacells and 69 Leiden clusters, with both levels of clustering containing a mixture of cells from the reference atlas^37^ and the present study. The most abundant cell type from the reference atlas in each Leiden cluster was used to assign preliminary labels to them and all the metacells therein.

Then, we filtered out the cells from the reference atlas and constructed a new set of metacells derived exclusively from the five time-points in the new dataset, using the same procedure outlined above. For each new metacell (297 in total) we retained the cell type annotations obtained from the above procedure, with the exception of the the splitting of the gland neurosecretory clusters into seven subtypes (instead of the original three; see below and Extended Data Fig. 1g). For each metacell and cell type, we calculated normalised gene expression values using the regu-larised geometric mean of gene counts for each cell group divided by the median across all clusters, as described in the *metacell* package^41^.

We validated the correspondence between cell type annotations in the new and reference datasets using *AUCell*^90^ 1.22.0. Specifically, we constructed a transcriptomic profile for each cell type in the reference atlas (consisting of a list of genes with a normalised fold change ≥ 2 in that cell type, up to 1,000 per cell type), constructed a ranking of the top genes per metacell and cell type in the new atlas (constructed with *AUCell::AUCell_buildRankings* and using the normalised fold change matrix as input), and scored the similarity of the reference profiles and the query rankings using *AUCell::AUCell_calcAUC* (restricting the search to the top 100 genes in each query cell type or metacell, using the *aucMaxRank* parameter). The resulting AUCell scores are scaled 0 to 1 and can be interpreted as the probability of finding genes from the reference cell type-specific profiles in the query rankings of top genes per cell type. Both the metacell and cell type matrices were visualised with the *ComplexHeatmap* 2.1.8 *R* library^91^ (Extended Data Fig. 1g).

#### Metacell tree reconstruction

We constructed hierarchical tree representations of the metacell clusters using *Bonsai*^92^ version 1.0.0. Specifically, we normalised the expression values of the metacell-level UMI count matrices using the *Sanity*^93^ 1.1 Bayesian framework, with default values of variance for the log transcription quotients (minimum of 0.001 and maximum of 50); and used these normalised values to build a tree representation using *Bonsai*, with a lenient cut-off for noisy genes (parameter *--zscore_cutofi 1.0*) and default parameters for nearest-neighbour tree space exploration (parameters *--nnn_n_ran-dommoves 1000*, *--nnn_n_randomtrees 20*, and *--use_knn 10*).

For visualisation purposes, we collapsed internal branches of the resulting tree based on the number of node-specific marker genes. Specifically, for each internal node in the tree, we selected all cells assigned to its daughter metacells, and calculated the number of marker genes with a fold-change value ≥ 1.5 and Bonferroni-adjusted *p*-value < 0.001 (Wilcoxon rank-sum tests on normalised counts), using cells descending from its sister node as background. This set of genes was further restricted to exclude genes that did not have at least a normalised fold-change value ≥ 1.5 in ≥ 70% of the daughter metacells, or had a normalised fold-change value ≥ 1.1 in ≥ 70% of the sister metacells. Finally, internal nodes with less than specific 10 markers were collapsed.

#### Condition stratification in metacell clusters

We evaluated the degree of stratification by time-course condition across metacell clusters by calculating the normalised entropy of the fraction of cells from each condition in groups of metacells defined from the *Bonsai* tree structure (Fig. 1e), taking the median value from all daughter metacells for each internal node. The normalised entropy statistic *e* ranges from *e =* 0 when each metacell is entirely composed of cells from one condition, to *e* = 1 when conditions are completely mixed within a metacell. The normalised entropy statistic was calculated using the *entropy* function in the *posterior* 1.7 R library^94^.

#### Batch efiects in the cell type clusters

We evaluated the compositional biases due to batch effects (i.e. each of the two 10x sequencing experiments) in the cell type clusters using two different procedures (Extended Data Fig. 1h). First, we used the *χ^2^* test implemented in *stats::chisq.test* to evaluate deviations from the expected proportions (48.7% of the cells from the first replicate, 51.3% from the second one) in each cell type. This naïve method revealed no significant deviations at *p* < 0.05 (Extended Data Fig. 1h); however, it does not account for correlated changes in the proportions of a given cell type due to changes in others, among other biases. Thus, we also used the *scCODA* Bayesian model^95^ implemented in the Python *pertpy* 0.10.0 library^96^ to test for compositional biases, using the cnidocyte cluster as a reference (selected because it did not exhibit changes in the *χ^2^* test analysis). At a false-discovery rate of 0.05, this model did not identify any batch effects either (the probabilities of inclusion of all cell types in the perturbed category was ≈ 0.5; Extended Data Fig. 1h).

#### Gene module inference and annotation

We used the *WGCNA* 1.73 *R* library^42^ to identify modules of co-expressed genes in our dataset. In order to better capture genes that co-vary according to the heat stress condition of the experiment (rather than purely due to cell type-level differences in gene expression), we constructed five dedicated sets of metacell clusters by re-clustering the cells from each condition (from control *t* = 0 h to recovery *t* = 52 days) and concatenating these sets of metacells. Each of the resulting 294 new metacell clusters, which contained only cells from the same condition (55 metacells for *t*= 0 h, 65 for *t* = 6 h, 67 for *t* = 48 h, 47 for *t* = 7 days, and 59 for *t* = 52 days), was annotated according to the majority cell type according to the labels drawn from the whole-atlas dataset.

We used the matrix of normalised fold-change gene expression values in this new set of metacells as input for *WGCNA*. Specifically, (i) we selected variable genes with a minimum fold change ≥ 1.2 in more than one metacell; (ii) we built a gene-gene co-expression matrix of Pearson correlation coefficients with *WGCNA::adjacency* (elevated to *power = 4*, a soft power parameter determined with *WGCNA::pickSoftThreshold* to test values between 2 and 20 and evaluating the fit of the scale-free topology model at each step); (iii) we used a hierarchical clustering approach to define gene modules using the *dynamicTreeCut::cutreeHybrid* function^97^ (parameters: *split* = 2, *minClusterSize* = 10, *cutHeight* = 0.99); (iv) we assigned each gene to one module (non-overlapping memberships) by applying a correlation threshold ≥ 0.5; and (v) we calculated the eigenvectors of each module across the input metacells using the *WGCNA::moduleEigengenes* function.

#### Diffierential gene expression analysis

We identified genes differentially expressed between sets of cells (selected according to specific stress conditions, cell types, metacell, etc., ass appropriate) with the *Seurat::FindMarkers* function. Specifically, we performed Wilcoxon rank-sum tests comparing normalised expression values in the two cell sets, and adjusting the *p*-values with the Bonferroni procedure. We did not filter out genes with low expression or low expression fold change values (parameters *min.pct = 0.0* and *logfc.threshold = 0*).

#### Re-clustering analysis of the gastrodermis

In order to gain a more granular insight into the cell type diversity of the gastrodermal cells, we performed a re-clustering analysis of all single-cell transcriptomes annotated as gastrodermis, gastrodermis-like, and alga-hosting (6,452 cells in total). For this purpose, we used the *metacell* algorithm (same procedure and parameters as for the whole-organism analysis, see above) to construct a total of 100 new metacells, which were annotated according to their majority cell type and stress stage in the whole-organism 10x dataset. The normalised fold-change gene expression values across these metacells were used to construct dedicated gastrodermis-specific gene modules with *WGCNA*, using the same procedure outlined above.

#### Analysis of the validated alga-positive single-cell transcriptomes

To map and quantify the MARS-seq transcriptomic data, we used a joint genome reference of *S. pistillata* and *Symbiodinium microadriaticum* (Symbiodiniaceae clade A, the dominant algal symbiont in *S. pistillata*). We mapped reads onto both genomes using *STAR*^98^ (with parameters *--outFilterMultimapNmax 1*, *--outFilterMismatchNmax 8* and *--alignIntronMax 6500*), applying downstream filters to quantify UMIs previously described^82^, (i.e. elimination of spurious UMIs resulting from synthesis and sequencing errors, elimination of artifacts involving unlikely IVT product distributions that are consequence of second strand synthesis or IVT errors). We used a minimum *FDR q*-value = 0.02 for quality filtering. The same procedure was repeated against a joint reference genome consisting of four Symbiodiniaceae species from four different clades: *Sy. microadriaticum* (clade A), *Breviolum minutum* (B), *Cladocopium goreaui* (C), and *Fugacium kawagutii* (F). The genome assembly and annotation versions used in this study are available in Supplementary Table 7.

Upon examination of the UMI/cell distributions against the coral and symbiont concatenated genomes (Extended Data Fig. 4a-c), we confirmed *Sy. microadriaticum* as the dominant symbiont in our dataset, and selected for further analysis profiles with at least 100 UMIs and 50 genes from *S. pistillata*, and at least 20 UMIs from *Sy. microadriaticum* (total *n* = 1,398, including 329, 98, 226, 339 and 406 cells from each condition from control to recovery). The UMI/cell recovery for the algal transcriptome was much lower than that of the host cell, thus precluding further analysis (Extended Data Fig. 4a). From this point on, these transcriptomic profiles were analysed using only counts from *S. pistillata* genes.

We constructed high-granularity metacell clusters for the alga-positive host cell transcriptomes using the same procedure outlined above. After removal of poorly supported cell clusters (with less than 80 specific gene markers or three transcription factors with normalised fold-change ≥ 2), we retained 1,082 cells, classified into 14 metacells, which covered all the time-course conditions (with 242, 93, 140, 287 and 320 cells from each condition from control to recovery, respectively). We built hierarchical tree representations of these metacells with *Sanity* and *Bonsai* (see above), and identified three main metacell clusters (canonical host cells, stressed host cells and recovery host cells; Fig. 4) based on node-specific gene markers (same procedure as described for the general atlas, but collapsing nodes with less than 5 specific markers and relaxing the Wilcoxon test stringency to *p*-value < 0.01). We also constructed modules of co-expressed genes using *WGCNA* (same procedure as above, except for an initial threshold of metacell-level normalised fold-change ≥ 1.5).

In order to cross-validate the existence of the alga-positive cell populations identified in the MARS-seq dataset (the main clusters plus the putative alga-engulfing cell population, represented by a single metacell; Fig. 4e), we used *AUCell* to compare the transcriptomic profiles of each of these sub-populations against all gastrodermal cells identified in Levy *et al.*^37^. Specifically, we selected marker genes from each cell population (those with normalised fold-change ≥ 1.5 at least 50% of their constituting metacells, and < 1.1 in 50% of the others) to build cell cluster profiles, and scored the similarity of these profiles against gastrodermal single-cell transcriptomes from both studies (using UMI count matrices as input for *AUCell::AUCell_buildRankings*), using using *AU-Cell::AUCell_calcAUC* (with *aucMaxRank = 1000*). For each cell population, we selected the best-fitting *AUCell* score threshold with *AUCell::AUCell_exploreThresholds* (parameters: *thrP = 1e-3, smallestPopPercent = 0.01*), and visualised the resulting histograms (*assignCells = TRUE*; Extended Data Fig. 4e).

#### Evolutionary conservation of gene expression programmes

We measured the level of evolutionary conservation of specific gene expression programmes (e.g. the set of stress-responsive genes in a given cell type, or gene modules), using a number of the genomic and transcriptomic conservation measurements (details below). In both cases, we compared the distribution of evolutionary conservation measurements between the gene set of interest (e.g. stress-responsive genes in a given cell population) against a matched background (other genes expressed in that cell population), using the log_2_ fold-change of the median values and two-sided Wilcoxon rank sum tests to assess the statistical significance.

For genomic comparisons, we calculated average *PhastCons* scores from various regions in each gene set (coding regions and *cis*-regulatory regions, see details below).

For transcriptomic comparisons, we used iterative comparisons of co-expression (*ICC*), from which we derived expression conservation scores as previously described^43,81^. Briefly, for a pair of species *a* and *b*, we concatenated their metacell-level normalised fold-change expression matrices, matched using one-to-one orthologous gene pairs (see below). The resulting matrices have *n* matched genes across *m_a_* and *m_b_* states (metacells). Then, we calculated the Pearson correlation of these two matrices, resulting in a *n×n* matrix, from where we can extract a vector of correlation values between orthologous pairs of genes from the diagonal. Then, the correlation vector (length *n*) is transformed into an initial weight vector (or initial expression conservation vector, *EC_0_*) by setting negative values to 0 (i.e. transforming from range −1 to +1, to 0 to 1). This weight vector is used to recalculate the *n×n* matrix using weighted Pearson correlation (instead of unweighted, as in the initial iteration) to produce a new weight vector (*EC_1_*). This procedure is repeated iteratively until the weights in iteration *i* and *i − 1* converge, i.e. *∑(EC_i_ − EC_i-1_)^2^* < 0.05. The final *EC* weight vector gives us the expression conservation scores between each pair of orthologous gene pairs. Expression conservation scores were calculated from metacell-level normalised fold-change matrices between *S. pistillata* and the following species: two scleractinian corals (*Oculina patagonica* and *Acropora millepora*^37^), a sea anemone (*N. vectensis*^99^) and a soft coral (*Xenia* spp.^35^). All the atlases used in these analyses were obtained from non-stressed specimens (including *S. pistillata*^37^).

### Comparative genomic analyses and functional gene annotation

#### Orthology and functional annotation of genes

We used the gene orthology databases for metazoans (24 species) and anthozoans (16 species) published in a previous study^37^. In short, these were produced using a combined approach based on (i) proteome-wide orthology assignments with *Broccoli*^100^ 1.2 and (ii) targeted phylogenetic analysis of transcription factor gene families, defined according to the detection of a curated set of Hidden Markov Model profiles (HMM) of known DNA-binding protein domains from the *Pfam* database^101^. All transcription factors from each family were retrieved using the *hmmsearch* utility in *HMMER*^102^ 3.3.2, aligned using *mafit*^103^ 7.47, gene trees obtained using *IQ-TREE*^104^ 2.1.0, and orthology groups and pairs are obtained using *Possvm*^105^ 1.1.0. The list of species with data sources is available in Supplementary Table 7.

Then, we annotated the genes in *S. pistillata* using the following sources of information: (i) gene names obtained from the *M. musculus* orthologs of each gene; (ii) Gene Ontology (GO) terms from their orthologs in *M. musculus*, as annotated in the November 2022 release of the Mouse Genome Database^106^; (iii) PFAM domain annotations for the predicted peptides of the longest isoform per gene, obtained with *Pfamscan* against release 33.1 of the Pfam database^101^.

#### Gene phylogenetic analysis

For the phylogenetic analysis of the *Xenia* sp. *LePin* lectin gene, we retrieved all peptide sequences from its orthologs from the Anthozoa orthology database (see above), constructed a *mafit* multiple sequence alignment of (*E-INS-i* algorithm with up to 10,000 refinement iterations), pruned the alignments with *clipkit*^107^ 1.1.395 (in *kpic-gappy* mode and using a gap threshold = 0.7), and built the gene tree using *IQ-TREE* 2.1.0 (up to 10,000 iterations until the convergence threshold = 0.999 is met for at least 200 generations; the best substitution model among LG, WAG and JTT was selected using *ModelFinder*^108^; statistical supports were calculated with 1,000 iterations of UFBoot^109^). Transmembrane domains were annotated using *TMHMM*^110^ 2.0, and signal peptides with *SignalP*^111^ 5.0b.

#### Functional gene set enrichment analysis

We performed functional enrichment tests of GO terms using the *topGO* 2.54.0 *R* library^112^. We computed the enrichments using counts of genes belonging to each relevant category (e.g. enriched markers in a cell type, gene module members, etc.) relative to all annotated and expressed genes, using Fisher’s Exact test and the *elim* algorithm for weighting of the GO ontology graph. For functional enrichment tests of Pfam domains, we used hypergeometric tests with *stats::phyper* (parameter *lower.tail = FALSE* for one-sided testing), comparing the frequencies of unique Pfam domains in each gene set of interest to their frequency in the background gene set.

#### Whole-genome alignments and sequence conservation analyses

We calculated genome-wide conservation scores *S. pistillata* using whole-genome alignments and *PHAST* (Phylogenetic Analysis with Space/Time) models. We built all-to-all whole-genome aligments of eight Pocilloporidae genomes using *Cactus*^113^ 3.0.1, guided by the species trees (Supplementary Table 7), and used *hal2maf* from the *HAL* 2.2 toolkit^114^ to build MAF alignments using scaffolds in *S. pistillata* as reference.

To identify conserved regions in the *S. pistillata* genome, we used *HAL* toolkit implementations of *Phast*^115^ utilities, as follows. First, we identified ancestral repetitive regions in *S. pistillata*, identified as annotated repetitive elements (annotated using EDTA^116^ with the *--anno 1* and *--cds* flags) with coordinates that could be lifted to the reconstructed Pocilloporidae last common ancestor’s genome (using *halLiftover*), and were longer than 100 bp in length in the extant reference genome. Second, we used these regions to train a null model of neutral substitutions in Pocilloporidae using *phyloFit*^117^ and the *halPhyloPTrain* utility, using the species tree indicated above, the *SSREV* substitution model, and the *--precision HIGH* and *--no4d* flags. Third, we used *PhastCons* to optimise this model using the expectation-maximisation procedure, re-estimating the transition probabilities and tree parameters at each iteration (flags *--target-coverage 0.25 --expected-length 12 --estimate-trees* and --*no-post-probs*). Finally, we calculated *PhastCons* scores for individual bases in the reference genome, as well as averages over coding and promoter regions (transcription start site −800 bp and +200 bp) for each gene. All calculations involving genomic coordinates were performed using *bedtools*^118^ 2.30 and the *R libraries GenomicRanges* 1.54, *IRanges*^119^ 2.36, and *rtracklayer*^120^ 1.62 *R* libraries.

### Code and library versions

All code for this project has been run on *R*^121^ version 4.3.1 and *Python*^122^ version 3.10, using the specific libraries mentioned in the Methods section. The code necessary to reproduce this study is available in Github: https://github.com/sebepedroslab/stylophora-heat-stress-sc-atlas.

## Author contributions

S.L. and A.S.-P. conceived the project. X.G.-B., L.M.-E., E.K., S.R.N., S.L., and A.S.-P. were involved in conducting the investigation, including experiments and data collection. X.G.-B. and S.L. were involved in formal analysis and creation of visualisations. X.G.-B. wrote the software code used in the analyses. T.M. and S.L. obtained the materials and resources. S.L. and A.S.-P. were in charge of funding acquisition, developing the methodology, and supervising this project. X.G.-B., S.L. and A.S.-P. wrote the original draft of the manuscript, and all the authors have read, reviewed and approved the final manuscript.

## Acknowledgements

Research in the A.S.-P. group has been funded by the European Research Council (ERC) under the European Union’s Horizon 2020 research and innovation programme (grant agreement number 851647), the BBVA Foundation (Ayudas Fundacion BBVA a Proyectos de Investigacion Cienti-fica 2021Proyecto CoralCellSeq), and the Spanish Ministry of Science, Innovation and Universities (MCIU; PID2021-124757NB-I00 funded by MCIN/AEI/10.13039/501100011033/FEDER, UE). S.L.’s group acknowledges support from the Israel Science Foundation (ISF grant number 2729/25). This project has received funding from two Marie Skłodowska-Curie grants from the EU Horizon 2020 programme (number 101065294, to S.L.; and 101031767, to X.G.-B.); from La Caixa Foundation (grant 100010434 to X.G.-B.; LCF/BQ/PR24/12050023); and from MCIU (grant PRE2022-105558, to L.M.-E.). We acknowledge support of MCIU through the Centro de Excelencia Severo Ochoa scheme (CEX2020-001049-S, MCIN/AEI/10.13039/501100011033), and from Generalitat de Catalunya through the CERCA programme. We thank the CRG Core Technologies Programme for their support and assistance.

## Extended Data Figure legends

**Extended Data Figure 1. Single-cell transcriptomic atlas quality control. a,** Saturation analysis for the two single-cell RNA-seq 10x libraries (replicates), measured as the fraction of genes identified genes at down-sampled sequencing depths. **b,** Distribution of detected unique molecule identifiers (UMI) per cell in each replicate. **c,** Normalised ClickTag (CT) counts as fraction of UMIs (rows) for each cell (columns). Each condition in the transcriptomic time-course has been labelled with three CT barcodes (colour-coded rows), and the normalised CT counts of the top three barcodes per cell is used to assign its single-cell RNA-seq profile to a specific condition (see Methods). **d,** Distribution of log_2_ ratios between the normalised counts of the top-three CT barcode combinations per cell, and the next most-abundant barcode combination. Cells are divided into seven categories: if the three top CT barcodes are assigned to any of the five time-course conditions, they are labelled accordingly; if not, they are classified as discordant or low-CT UMI cells (see Methods). **e,** Joint two-dimensional uniform manifold approximation projection (UMAP) of the *S. pistillata* transcriptomic time-course generated in this study (gray outline) and the previous data-set (Levy et al. 2025), from which the cell type labels (colour-coded dots) have been lifted. **f,** Fraction of cells originating from each of the two replicates in this study and the Levy *et al.* 2025 study, for each of the cell types annotated therein. **g,** Transcriptomic similarity between the cell types annotated in Levy *et al.* 2025 (columns) and metacells (rows; top panel) or cell types (rows; bottom panel) in this study, measured as *AUCell* scores. **h,** Fraction of cells from each replicate in the cell types annotated in this study (barplot on the left); *p-*values from two-sided *χ^2^* tests (middle) measuring the deviation from the global expected proportions (48.7% from replicate 1); and inclusion probability (IP) and fold-change (FC) values obtained from *scCODA* tests of cell type composition perturbation (right). **i,** Fraction of UMIs per gene in each of the biological replicates for each condition in the time-course, and correlation coefficient (*r*) and *p*-value from Pearson’s product-moment correlation tests between replicates.

**Extended Data Figure 2. Gene module analysis. a,** Heatmap showing the level of over- or un-der-expression of modules of co-expressed genes (rows, colour-coded by the module expression pattern) in specific cell types, comparing each condition to the control *t* = 0 h condition (columns, colour-coded by cell type and condition). Fold-change values (colour-coded in the heatmap) are defined as the ratio of the median fraction of UMIs from a module in a given cell type and condition, compared to *t* = 0 h in that cell type. Statistical significance (*p-*values, denoted by dot size) was assessed using Wilcoxon rank-sum two-sided tests (available in Supplementary Table 3). **b,** Distribution of UMI fractions per cell for five stress-related gene modules, in selected cell types where they are active and variable. The statistical significance of differences to *t* = 0 h was assessed using Wilcoxon rank-sum two-sided tests (exact *p-*values shown). **c,** Distribution of UMI fractions per cell for the innate immunity module in three cell types (epidermis, gastrodermis and *Pou4+* neurons), stratified by time-course condition and by biological replicate.

**Extended Data Figure 3. Per-cell type differential gene expression analysis. a,** Evolutionary conservation of genomic (left) and transcriptomic features (right) of the gene sets differentially expressed during the stress/recovery time-course in each cell type or population (rows). Genomic conservation for the coding sequence (CDS) and promoter regions of each gene was measured using median *Phastcons* scores; transcriptomic conservation in other species (the scleractinians *O. patagonica* and *A. millepora*, the sea anemone *N. vectensis*, and the soft coral *Xenia sp*.) was measured using expression conservation scores derived from iterative comparisons of gene co-expression (ICC) between non-stressed single-cell transcriptomic atlases, at the metacell level (see Methods). Fold-changes (colour-coded in the heatmap) are calculated from comparing the median values of the genes of interest and the genomic background (other expressed genes); statistical significance (*p-*values, denoted by dot size) was assessed using Wilcoxon rank-sum two-sided tests. **b,** Distribution of gene ages (phylostrata) for the stress-responsive gene sets (all cell types), compared to a genomic background of non-stress-responsive expressed genes (top); and log_2_ odds-ratio of enrichment of each phylostrata (*p-*value from two-sided Fisher’s exact tests; bottom). **c,** Evolutionary conservation of genomic (left) and transcriptomic features (right) of the gene sets associated to specific gene modules (fold-changes and *p*-values calculated as in panel **a**). **d,** Gene age analysis of stress gene modules (log_2_ odds-ratio of enrichment of each phylostrata, *p-*values as in panel **b**). **e,** Enrichment analysis for the membership to specific gene modules (col-our-coded bars) among all genes differentially expressed in a given cell type during the stress and recovery time-course (adjusted *p*-value < 0.001 and absolute fold-change > 1.1). For each gene module, we show the log_2_ of the odds ratio of Fisher’s exact test (vertical axis), and the number of genes mapping to that module and the significance of the enrichment (dots and numbers below the barplots, colour-coded by enrichment *p-*value). Modules are colour-coded according to their transcriptional profile and inferred function, with transversal modules in gray (with dark gray indicating stress-related modules) and cell type-specific modules coloured accordingly.

**Extended Data Figure 4. Cell population dynamics of alga-hosting cells. a,** Number of UMIs/cell detected in each MARS-seq transcriptome, for *S. pistillata* genes (horizontal axis) and *Symbiodinium microadriaticum* genes (vertical axis). Dotted lines indicate the UMI/cell thresholds used to filter out low-UMI single-cell transcriptomic profiles (100 UMI/cell). The adjacent histograms show the distribution of UMIs/cell in each species. **b,** Overlap of the sets of cells in with ≥ 100 UMI/cells from *S. pistillata* and *Sy. microadriaticum* in the MARS-seq transcriptomic atlas. **c,** Number of UMIs/cell detected in the MARS-seq transcriptome for *Sy. microadriaticum* genes (vertical axis) and three other Symbiodiniaceae species, representative of clades B (*Breviolum minutum*), C (*Cladocopium goreaui*) and F (*Fugacium kawagutii*). **d,** Distribution of UMI/cell from *S. pistillata* (top) and *Symbiodinium microadriaticum* (bottom) genes in each metacell, with the metacell tree shown for reference. **e,** Distribution of *AUCell* similarity scores obtained from scoring the transcriptomic profiles of specific cell populations identified in the alga-positive MARS-seq time-course (canonical host cells, stressed and recovering host cells, and the putative alga-engulfing cell population; Fig. 4e) against the gastrodermal cell transcriptomes from Levy *et al.* 2025, with the number and percentage of signature-positive cells indicated in each case. Dotted vertical lines indicate the *AUCell* score thresholds used to identify signature-positive cells in each case (using a maximum stringency criterion). Barplots next to each histogram indicate the fraction of cell type annotations in the signature-positive cells. **f,** Heatmap of expression of differentially expressed genes in non-host, alga-engulfing, and canonical alga-hosting cells (shown as gene-scaled UMI fractions in each cell population). Insets to the right indicate the fold-change (colour-coded) and statistical significance of differential expression (Bonferroni-adjusted *p-*values from Wilcoxon rank-sum tests on normalised counts; shown by dot size) for selected genes relative to the control condition. Genes are colour-coded according to selected gene module membership. The table to the right indicates whether selected C-lectin genes are orthologs of the *LePin* gene of *Xenia* sp., and whether they have transmembrane domains or secretion signal peptide (see panel g). **g,** Phylogenetic tree of *Xenia* sp. *LePin* gene (Xesp_002591-T1) orthologs in other anthozoan species, colour-coded in the three tips by taxonomy. The table to the right indicates whether C-lectin genes from *S. pistillata* are significantly over-expressed in the alga-engulfing cell population (Bon-ferroni-adjusted *p-*value < 0.001 and fold-change > 1.1), and whether they have transmembrane domains or secretion signal peptides, and their PFAM domain architectures. Sequence features are also shown for other selected species. The complete tree and the alignment is available in Supplementary Table 7.

## Supplementary Table legends

**Supplementary Table 1. Counts of Symbiodiniaceae cells in the coral nubbins. a,** Cell counter measurements of Symbiodiniaceae cells (based on chlorophyll autofluorescence) across five conditions in the stress time-course (*t* = 0 h, 6 h, 48 h, 7 d, and recovery). Measurements include three biological replicates (A to C), two nubbins per biological replicate, and three measurements per nubbin. In each instance, we report the total cell concentration, the concentration of chlorophyll-positive (CY5+) cells, and the corresponding percentage of chlorophyll-positive cells (Symbiodiniaceae symbionts). **b,** Per-nubbin mean fraction of chlorophyll-positive cells, and standard deviations (calculated from repeated measurements). **c,** Per-biological replicate mean fraction of chlorophyll-positive cells (calculated from fragment averages). **d,** Per-condition mean percentage of chlorophyll-positive cells, standard deviations, standard error of the mean, and ranges (from biological replicates). **e,** Results from a one-way analysis of variance (*ANOVA*) based on biological replicate means (top), and a repeated-measures *ANOVA* based on biological replicate means keeping individual specimens as subjects of the analysis (middle), and pairwise comparisons between conditions using Welch two-sample *t*-tests (bottom), for which *p-*values have been adjusted for multiple comparisons using Benjamini-Hochberg’s *FDR* procedure.

**Supplementary Table 2. Photosynthetic efficiency cells of coral-associated Symbiodiniaceae in the coral nubbins. a,** Photosynthetic performance measurements (*F_v_/F_m_*) across five conditions in the stress time-course (*t* = 0 h, 6 h, 48 h, 7 d, and recovery). Measurements of three biological replicates (A to C), two nubbins per biological replicate, and two measurements per nubbin. **b,** Per-nubbin mean *F_v_/F_m_*, and standard deviations (calculated from repeated measurements). **c,** Per-biological replicate mean *F_v_/F_m_*, and standard deviations (calculated from nubbin averages). **d,** Per-condition mean *F_v_/F_m_*, standard deviations, standard error of the mean, and rages (calculated from biological means). **e,** Results from a one-way repeated-measures *ANOVA* based on biological replicate means across conditions (top), and pairwise comparisons between conditions using Welch two-sample *t*-tests (bottom), for which *p-*values have been adjusted for multiple comparisons using Benjamini-Hochberg’s *FDR* procedure.

**Supplementary Table 3. Gene module memberships and functions. a,** Gene module assignments for 9,696 variable genes. Each gene is assigned to one module, named based on their inferred function and expression profiles. We report the gene name (inferred from mouse orthologs) and PFAM domains. **b,** Functional enrichment analysis of each gene module (browsable from the first column). For each gene module, we list all genes annotated to significantly enriched Gene Ontology (GO) terms (at *p*-value < 0.05 in Fisher’s Exact tests with *elim* correction), the number of genes annotated to each significant GO term, the PFAM domain architecture of each gene, and the results from a PFAM domain enrichment analysis using hypergeometric tests (only domains significant at *p* < 0.05). Two generic cell component terms (cytosol and cytoplasm) have been excluded for clarity. **c,** Change in activity of each gene module along the transcriptomic time-course, measured as the log_2_ fold-change between the median fraction of UMIs from that module in a given cell type and condition, and the median in either the control (*t =* 0 h) or the previous time-point. Statistical significance has been tested with two-sided Wilcoxon rank-sum tests.

**Supplementary Table 4. Per-cell type differential gene expression analysis, using the control as reference. a,** Differentially expressed genes in various cell populations (browsable from the first column) each time-course condition, compared *t* = 0 h. For each gene, we report the log_2_ fold-change value of mean normalised counts between two conditions (e.g. *t* = 6 h to 0 h ratio), *p*-values from two-sided Wilcoxon rank-sum tests (raw and Bonferroni-adjusted per cell type and time-course comparison), the fraction of UMIs in both conditions, the fraction of cells where the gene is detected in both conditions, and the total number of UMIs in that cell type; as well as its PFAM domain, gene name inferred from mouse orthologs, and gene descriptions. The cell populations tested in this analysis include all cells, all individual cell types, and groups of cell types (e.g. all neuronal cells considered together, as “neurons, all cells”). **b,** Functional enrichment analysis of the sets of differentially expressed genes in each cell population (browsable from the first column), condition comparison (e.g. *t* = 6 h to 0 h; browsable from the second column), and direction of enrichment (e.g. upregulated in *t* = 0 h or *t* = 6 h, third column). For each cell type and comparison, we list all genes annotated to significantly enriched Gene Ontology (GO) terms (at *p*-value < 0.05 in Fisher’s Exact tests with *elim*correction), the number of genes annotated to each significant GO term, the PFAM domain architecture of each gene, and the results from a PFAM domain enrichment analysis using hypergeometric tests (only domains significant at *p* < 0.05). Two generic cell component terms (cytosol and cytoplasm) havimme been excluded for clarity.

**Supplementary Table 5. Per-cell type differential gene expression analysis, using the previous time-point as reference. a,** Differentially expressed genes in various cell populations (browsable from the first column) and sequential combinations of time-course conditions (i.e. each condition is compared to its previous time-point). Reported values as in Supplementary Table 4a. **b,** Functional enrichment analysis of the sets of differentially expressed genes in each cell population (browsable from the first column), condition comparison (e.g. *t* = 48 h to 6 h; browsable from the second column), and direction of enrichment (e.g. upregulated in *t* = 48 h or *t* = 6 h, third column). Reported values as in Supplementary Table 4b.

**Supplementary Table 6. Per-cell type differential gene expression analysis and functions. a,** List of genes assigned to the alga-hosting stressed module (*n* = 444 genes). We report the gene name (inferred from mouse orthologs) and PFAM domains of each gene. **b,** Gene markers identified in the alga-engulfing gastrodermis cells, identified by differential expression analysis between cells from metacells with active engulfment gene module (eigenvalue *Z* > 1.28) and the rest of the gastrodermis, or the canonical alga-hosting cells (browsable from the first column). For each gene, we report the log_2_ fold-change value of mean normalised counts between the two cell sets (ratio between the putative alga-hosting population to the other cells), *p*-values from two-sided Wilcoxon rank-sum tests (raw and Bonferroni-adjusted per comparison), the fraction of UMIs in both conditions, the fraction of cells where the gene is detected in both conditions, and the total number of UMIs; as well as its PFAM domain and gene name inferred from mouse orthologs. **c,** Functional enrichment analysis of the gene sets enriched in the stress module and the putative alga-engulfing cells (browsable from the first column). For each gene set, we list all genes annotated to significantly enriched Gene Ontology (GO) terms (at *p*-value < 0.05 in Fisher’s Exact tests with *elim* correction), the number of genes annotated to each significant GO term, the PFAM domain architecture of each gene, and the results from a PFAM domain enrichment analysis using hypergeometric tests (only domains significant at *p* < 0.05). Two generic cell component terms (cytosol and cytoplasm) have been excluded for clarity.

**Supplementary Table 7. Genome data sources.** Taxon sampling, source studies and accession codes for the genomic datasets used for comparative genomic analyses in this study: the Anthozoa and Metazoa datasets used to build the gene orthology databases, and the Pocilloporidae-specific dataset used in the whole-genome alignment analyses. It also includes the list of Symbiodiniaceae genomes used to map the single-cell transcriptomic experiments.

