## Supplementary figures and images for "Cell-resolved transcriptional responses during heat-induced coral bleaching and recovery"

### Extended Data Figure 1

**Extended Data Figure 1**

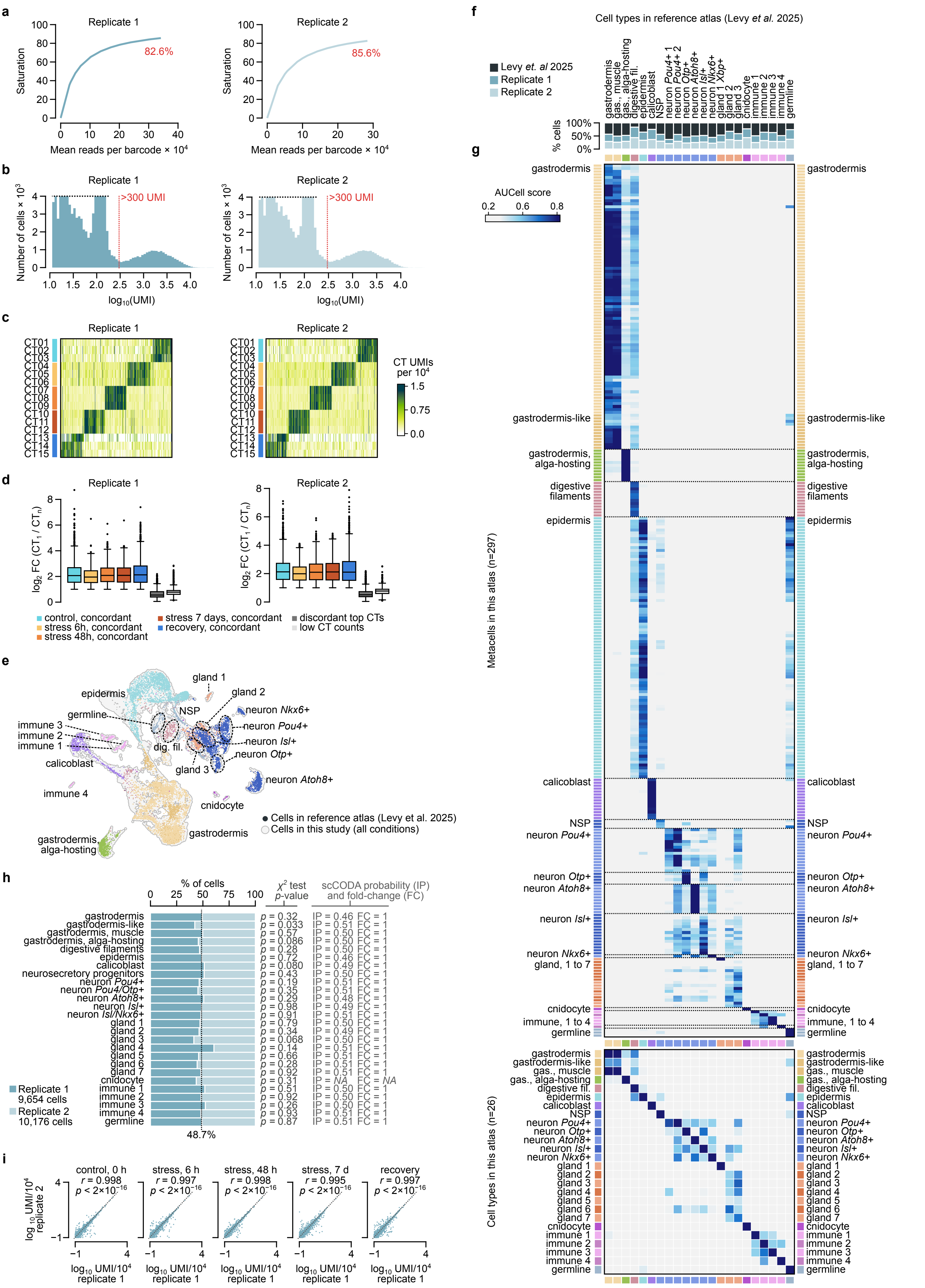

### Extended Data Figure 2

#### Extended Data Figure 2

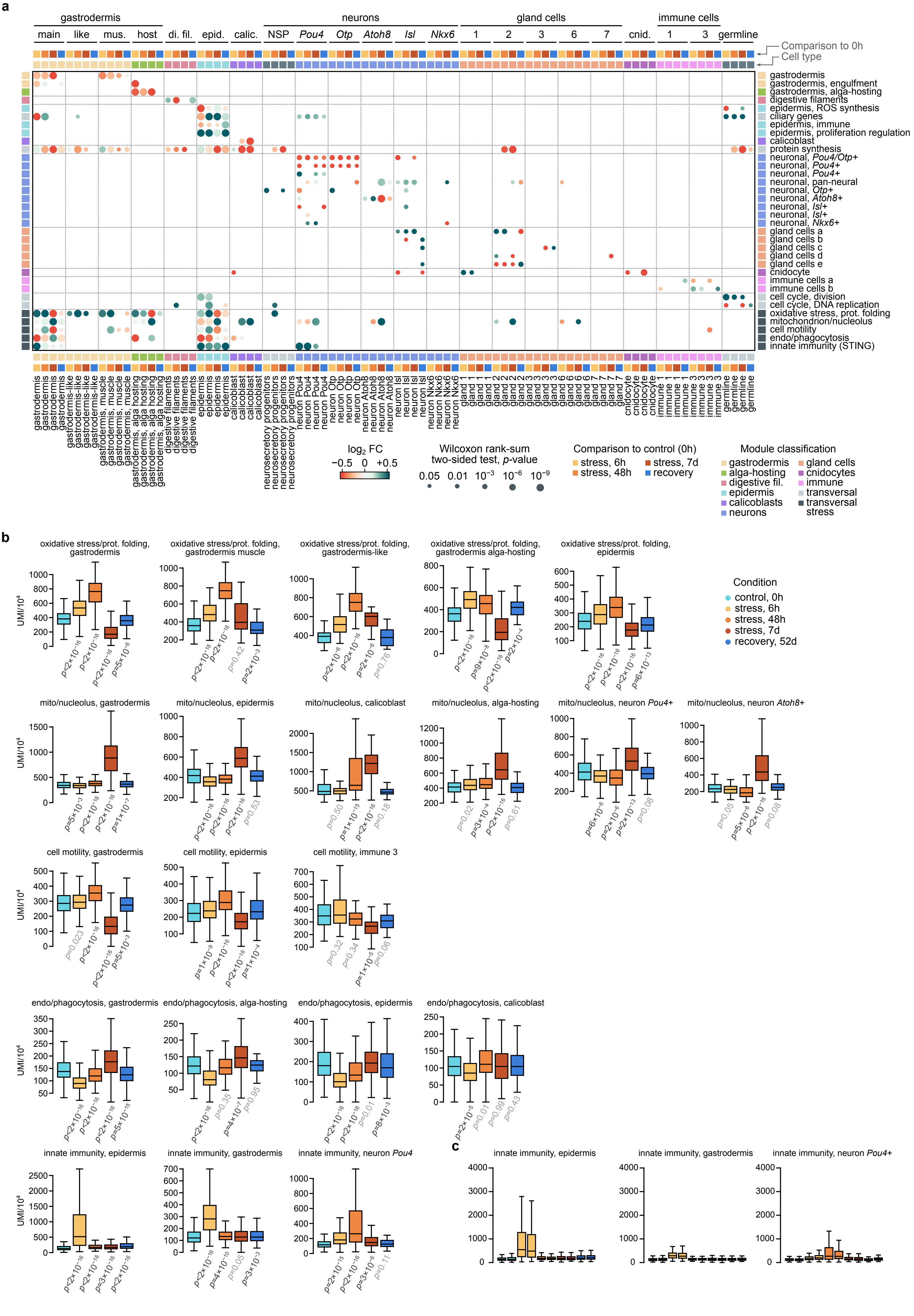

### Extended Data Figure 3

**Extended Data Figure 3**

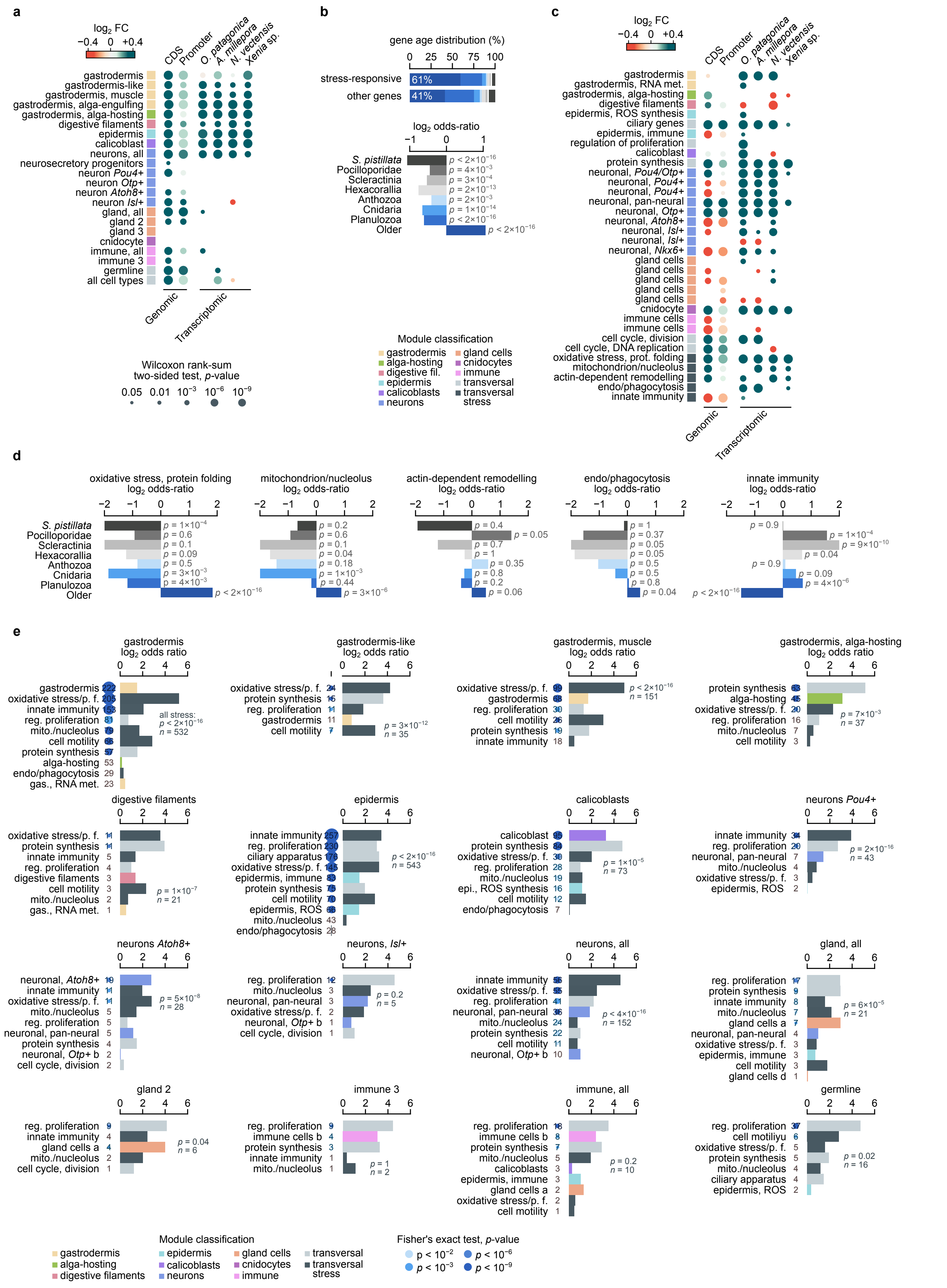

### Extended Data Figure 4

#### Extended Data Figure 4

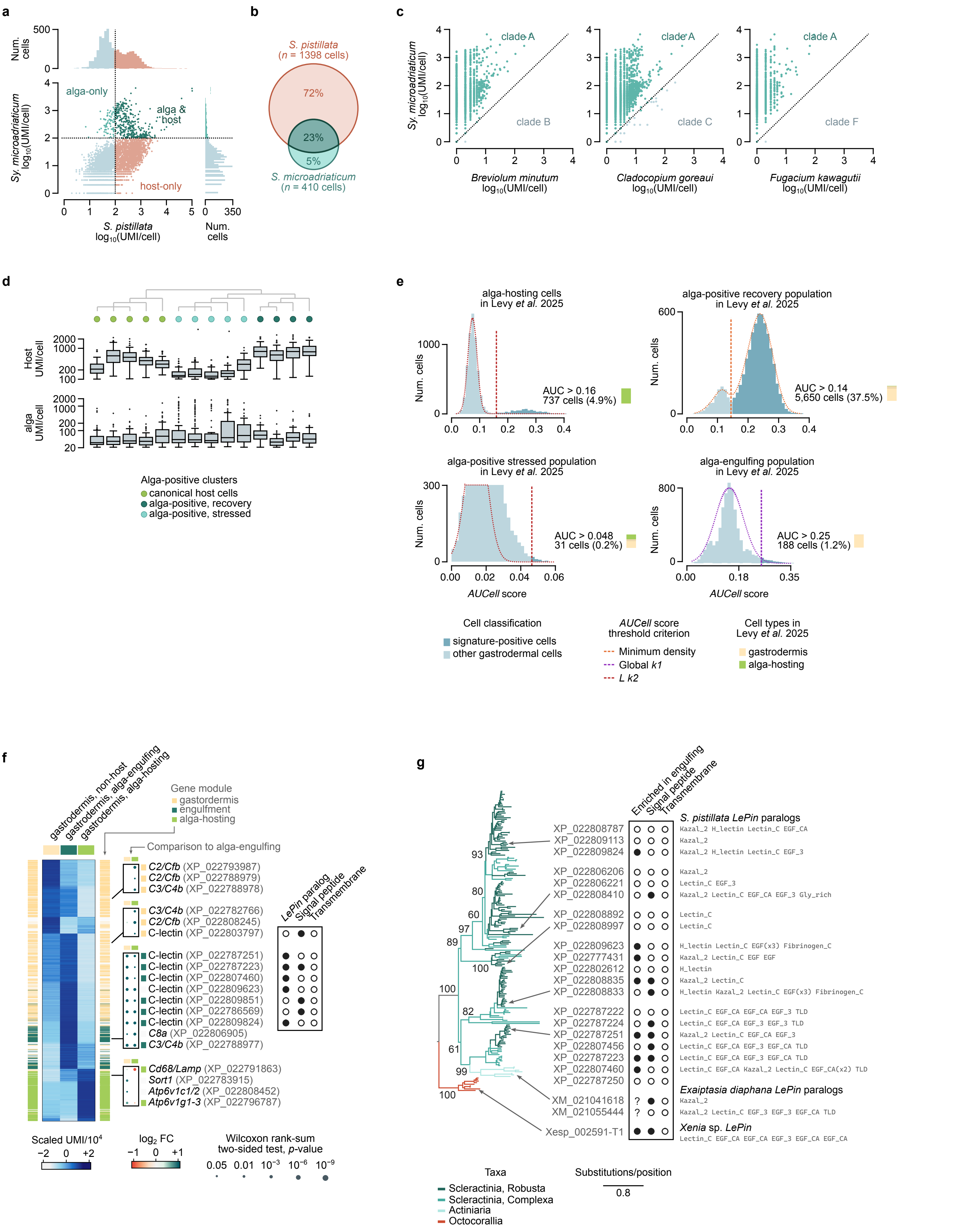
